# Six states of *Enterococcus hirae* V-type ATPase reveals non-uniform rotor rotation during turnover

**DOI:** 10.1101/2022.08.09.503272

**Authors:** Raymond N. Burton-Smith, Chihong Song, Hiroshi Ueno, Takeshi Murata, Ryota Iino, Kazuyoshi Murata

**Affiliations:** Exploratory Research Center on Life and Living Systems (ExCELLS), National Institute for Natural Sciences, Okazaki, Aichi, 444-8585, Japan; National Institute for Physiological Sciences, National Institute for Natural Sciences, Okazaki, Aichi, 444-8585, Japan; Department of Physiological Sciences, School of Life Science, The Graduate University for Advanced Studies (SOKENDAI), Okazaki, Aichi, 444-8585, Japan; Department of Applied Chemistry, Graduate School of Engineering, The University of Tokyo, 7-3-1 Hongo, Bunkyo-Ku, Tokyo 113-8656, Japan; Department of Chemistry, Graduate School of Science, Chiba University, 1-33 Yayoi-Cho, Inage-Ku, Chiba 263-8522, Japan; Institute for Molecular Science, National Institute for Natural Sciences, Okazaki, Aichi, 444-8787, Japan; Department of Functional Molecular Science, School of Physical Sciences, The Graduate University for Advanced Studies (SOKENDAI), Okazaki, Aichi, 444-8585, Japan

**Keywords:** V-ATPase, ATP-driven ion pump, off-axis rotor, membrane protein complex, cryo-electron microscopy, single particle analysis

## Abstract

The vacuolar-type ATPase from *Enterococcus hirae* (EhV-ATPase) is a thus-far unique adaptation of V-ATPases, as it performs Na^+^ transport and demonstrates an off-axis rotor assembly. Recent single molecular studies of the isolated V_1_ domain have indicated that there are subpauses within the three major states of the pseudo three-fold symmetric rotary enzyme. However, there was no structural evidence for these. Herein we activated the EhV-ATPase complex with ATP and identified multiple structures consisting of a total of six states of this complex by using cryo-electron microscopy. The orientations of the rotor complex during turnover, especially in the intermediates, were not as perfectly uniform as expected. The densities in the nucleotide binding pockets in the V_1_ domain indicated the different catalytic conditions for the six conformations. The off-axis rotor and its’ interactions with the stator a-subunit during rotation suggests that this non-uniform rotor rotation is performed through the entire complex.

## Introduction

Most rotary ATPases are a structurally similar but broad ability group of enzymes, which carry out a variety of different cellular processes (Stewart et al., 2014). They play critical roles within both prokaryotic and eukaryotic organisms, acting for ATP (adenosine triphosphate) synthesis (Kühlbrandt and Davies, 2016) and ATP-driven ion transport (Stewart et al., 2014). Their ubiquity, critical role in life of a cell, and sophisticated mechanism and design have made them popular targets for pharmacological treatments for a range of diseases from cancers (Martinez-Zaguilan et al., 1993), osteoporosis (Yuan et al., 2010), and kidney disease (Wagner et al., 2004) to bacterial and fungal infections (Eaton et al., 2021). They are classified by type; A-, F-, or V-type, which are not necessarily mutually exclusive in role (Grüber et al., 2001). For example, a single ATPase can carry out ATP synthesis as well as ion translocation by ATP hydrolysis (Weber and Senior, 2000).

The vacuolar-type (V-) ATPases are a ubiquitous class of membrane protein complex which utilise ATP hydrolysis to power ion transport across cellular membranes (Vasanthakumar and Rubinstein, 2020), although the *Thermus thermophilus* (Tt) V/A-ATPase is a dual-role ATPase (Yokoyama and Imamura, 2005), which usually functions as an ATP synthase (Nakano et al., 2008). The structures of the TtV/A-ATPase and *Saccharomyces cerevisiae* (Sc) V-ATPases and examinations of the subunits revealed common molecular mechanisms across V-ATPases (Schep et al., 2016).

V-ATPases are formed from two domains: the V_o_ and V_1_ domains. In the V-ATPase from *Enterococcus hirae* (EhV-ATPase), the V_1_ domain in cytoplasm consists of A-, B-, D-, and F-subunits, termed to be A_3_B_3_DF complex, and the V_o_ domain bound membrane consists of a-, c-, d-, E-, and G-subunits, termed to be ac_10_dE_2_G_2_ complex (Fig. 1). The cylindrical “c-ring” is located entirely within the membrane and binds the ion to be transported, rotating almost a full turn to bring the bound ions from ingress to egress. The a-subunit possesses a membrane-integrated region that interacts with the c-ring to control ion binding, and a cytosolic region that interacts with the EG “stalk” complexes to support the cytosolic V_1_ domain. The V_1_ domain creates a stator formed with the A_3_B_3_ complex connected to the EG stalks, and maintains the rotor shaft formed with the D- and F-subunits. The transport of the bound ion across the membrane is performed by the hydrolysis of ATP in the V_1_ domain, which drives the rotation of a coupling “rotor” at three pauses separated by 120°, depending on the pseudo trimeric A/B dimer V_1_ ATPase domain. This discontinuous rotation causes the rotation of the d-subunit and membrane-intrinsic c-ring, resulting in three conformational states in the entire complex. Many structures of V-ATPases have been reported demonstrating the three turnover states of each, both as isolated V_1_ domains (Arai et al., 2013; Maruyama et al., 2019; Suzuki et al., 2016) and as complete complexes (Abbas et al., 2020; Nakanishi et al., 2018; Wang et al., 2020a, 2020b; Zhao et al., 2015; Zhou and Sazanov, 2019).

**Figure 1.**
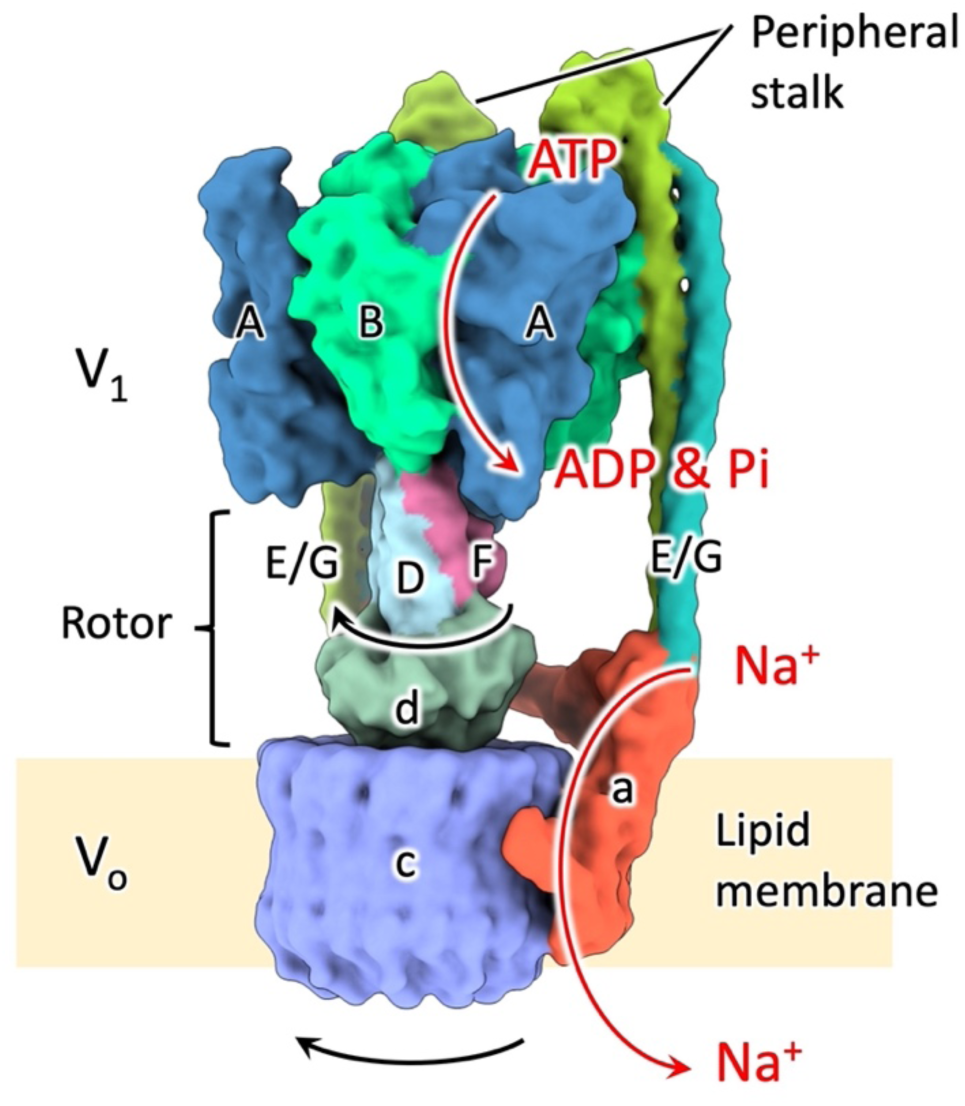
Schematic drawing of the EhV-ATPase. Subunits are individually color-coded and labelled. A, B, D and F-subunits form the V_1_ domain, while a-, c-, d-, E- and G-subunits form the V_o_ domain. The a-subunit and c-ring are embedded in lipid membrane. The rotation of the D/F/d rotor shaft and c-ring proceeds clockwise when viewed from top to bottom, as indicated. Turnover is driven by the entry of ATP into a binding pocket at the interface of each A/B dimer, the hydrolysis reaction of which drives conformational changes which cause the rotation of the rotor.

The V-ATPases show adaptations across species; the structures of the membrane-intrinsic c-ring and the cytosolic supporting peripheral stalks vary significantly in subunit composition and arrangement (Vasanthakumar and Rubinstein, 2020). For example, eukaryotic ScV-ATPase exhibits a large diameter c-ring consisting of 10 quadruple-helix subunits (a total of 40 transmembrane helices, although the subunits are not all homogeneous (Vasanthakumar et al., 2019)) and three rigid supporting stalks (Zhao et al., 2015), while prokaryotic TtV/A-ATPase exhibits a smaller diameter c-ring consisting of 12 twin-helix subunits (a total of 24 transmembrane helices) and just two supporting stalks (Nakanishi et al., 2018). F-ATPases use a single supporting stalk, which proves to be considerably more robust in form (Murphy et al., 2019).

*E. hirae* (formerly, *Streptococcus faecalis*) is a Gram-positive zoonotic bacterium with a sequenced genome with implication for both human health and the farming industry (Larsson. J. et al., 2014; Pinkes et al., 2019). It is a widely used model organism for bioenergetics, copper homeostasis, and ion transport studies (Ikegami et al., 1999; Kakinuma et al., 1999; Solioz and Stoyanov, 2003). The EhV-ATPase is specific for Na^+^ (Murata et al., 2000) rather than acting to control H^+^ gradient across the membrane, although Li^+^ transport has also been demonstrated (Furutani et al., 2011; Kakinuma et al., 1999; Kawano-Kawada et al., 2011). However, it has been reported that H^+^ can also be transported in severely Na^+^-depleted conditions (Leone et al., 2015; Murata et al., 2008). Subunit composition of the EhV-ATPase has been previously reported (Murata et al., 1997), being the same as the subunit composition of TtV/A-ATPase except for the number of subunits in the c-ring. As the structures of the isolated components of the complex, the atomic structure of the c-ring membrane-bound domain was elucidated by x-ray crystallography which demonstrated a large homogeneous decameric c-ring consisting of 10 quadruple-helix subunits (Murata et al., 2005). The structure illustrates that the ion selectivity is controlled by amino-acid residues in the c-subunit ion binding pocket (Kawano-Kawada et al., 2012). Furthermore, the complex structures of the rotor shaft (Saijo et al., 2011) and the pseudo-trimeric A/B dimer V_1_ ATPase domain (Arai et al., 2013) were determined by x-ray crystallography. The different catalytic states of the V_1_ domain were revealed using non-hydrolysable ATP analogues (Suzuki et al., 2016).

EhV-ATPase was previously reported to have no intermediate pauses between the 120° rotations of the DF rotor complex (Minagawa et al., 2013; Ueno et al., 2014). However, more recent work has demonstrated that there are indeed intermediate pauses divided by an approximately 40°/80° split between the 120° major pauses (Iida et al., 2019). Like the F-ATPases (Yasuda et al., 2001), which demonstrate subpauses between the distinct rotor positions aligned with the V_1_ domain, these subpauses are also not precisely midway between the major conformations. The F-ATPases show variability in angular subpauses, ranging from an 80°/40° split for thermophilic *Bacillus* PS3 (Masaike et al., 2008), a 65°/25°/30° split for human mitochondrial F-ATPase (Suzuki et al., 2014), an 85°/35° split for *Escherichia coli* (Bilyard et al., 2012), and an 87°/33° split for yeast mitochondrial F-ATPases (Steel et al., 2015). Recently, cryo-electron microscopy (cryo-EM) has visualized the subpause structures of the thermophilic *Bacillus* PS3 F_1_-ATPase (Sobti et al., 2021). However, there is still no structural evidence for subpauses in V-ATPases.

Previously, we reported the off-axis rotor (Tsunoda et al., 2018) of the EhV-ATPase using phase contrast cryo-EM with a Zernike-type phase plate. This permitted the first visualisation of the detergent-solubilized EhV-ATPase full complex. While this data provided insight into the overall macromolecular organisation of this unusual V-ATPase complex conformation, it was limited by resolution constraints. The off-axis nature of the rotor has thus far only been identified in EhV-ATPase (Tsunoda et al., 2018). Further, EhV-ATPase possesses a large diameter c-ring (Murata et al., 2005) akin to the eukaryotic form, while the V_1_ domain was supported by the two peripheral stalks, like other prokaryotic V-ATPases. The asymmetric stalks have been shown to play an important role in torque generation in EhV-ATPase (Ueno et al., 2014).

Here, we present six state structures of the EhV-ATPase complex in three major states corresponding to the three reported conformations in TtV/A-ATPase (Nakanishi et al., 2018) at resolutions of 4.2 Å to 4.4 Å, and three intermediate states at resolutions of 4.8 Å to 7.7 Å. Focussed refinements of the V_1_ domain improved the resolution to <4 Å in State 1. Importantly, these states were all identified using the same conditions and from the same preparation and grids, while many other works identifying intermediates utilise different conditions and substrates to promote the different conformations (Kishikawa et al., 2021). These results provide new insights into the V-ATPases, where the orientations of the rotor complex during turnover, especially in the intermediates, were not as perfectly uniform as expected. The off-axis rotor and its’ interactions with the stator a-subunit during rotation suggests that this non-uniform rotor rotation is performed through the entire complex.

## Results

### Data collection and 3D reconstruction of cryo-EM SPA for EhV-ATPase

To attempt to generate all catalytic states of EhV-ATPase, ATP, Mg^2+^, and Na^+^ were added to the purified and detergent solubilized complex. The sample was vitrified during the active state transition. A total of 47,372 micrograph movies were acquired using SerialEM (Mastronarde, 2005) on a JEOL CryoARM-300 electron microscope equipped with a Gatan K3 direct detector in three sessions (Table 1). The detergent solubilized EhV-ATPase was a very low contrast complex, which made it difficult to see in conventional defocus-acquired cryo-EM (Fig. S1A). As a result, the particle picking strategy used was more liberal, with a lot of undesirable particles in the selection of 4,383,273 particle candidates (Fig. S1B, Table 1). These particles were cleaned through multiple rounds of two-dimensional (2D) and three-dimensional (3D) classification. The processing strategy covered in detail in Methods and Fig. S1 broadly follows the standard processing workflow of RELION software (Fernandez-Leiro and Scheres, 2017; Scheres, 2012; Zivanov et al., 2018, 2020). Interestingly, without the addition of Na^+^, the purified EhV-ATPase would tear itself apart upon addition of ATP. Despite this, the complex was still very fragile, and a large percentage of particles were removed during classification due to evident damage to either the rotor shaft or the V_o_ domain (Fig. S1, Table 1). The ice-embedded complex showed a strong orientation preference (Fig. S2A), which limited the resolution and quality of reconstructions with the first acquired dataset. For this reason, the third dataset was acquired with both 20° and 30° stage tilt to attempt to overcome this orientation preference (Table 1). It was mostly successful (Fig. S2B), and provided sufficient extra variance in views to improve the angular sampling of reconstructions (Fig. S2C).

**Table 1.**
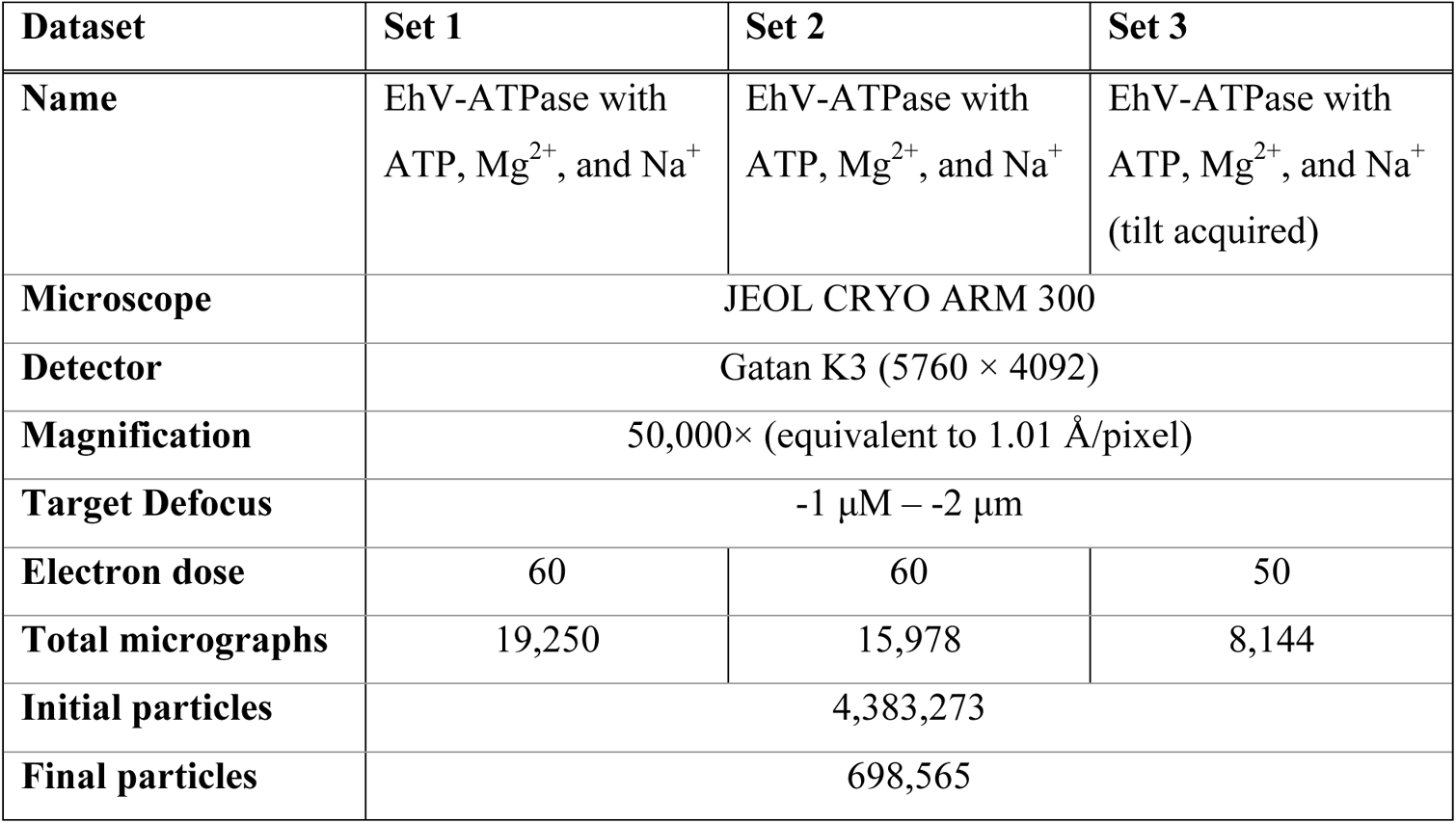
Data collection and image processing information for EhV-ATPase treated with ATP, Mg^2+^, and Na^+^.

| Dataset | Set 1 | Set 2 | Set 3 |
| --- | --- | --- | --- |
| Name | EhV-ATPase with ATP, $\text{Mg}^{2+}$, and $\text{Na}^+$ | EhV-ATPase with ATP, $\text{Mg}^{2+}$, and $\text{Na}^+$ | EhV-ATPase with ATP, $\text{Mg}^{2+}$, and $\text{Na}^+$ (tilt acquired) |
| Microscope | JEOL CRYO ARM 300 |  |  |
| Detector | Gatan K3 (5760 × 4092) |  |  |
| Magnification | 50,000× (equivalent to 1.01 Å/pixel) |  |  |
| Target Defocus | -1 $\mu\text{M}$ – -2 $\mu\text{m}$ | | |
| Electron dose | 60 | 60 | 50 |
| Total micrographs | 19,250 | 15,978 | 8,144 |
| Initial particles | 4,383,273 |  |  |
| Final particles | 698,565 |  |  |

Focussed refinement using a soft mask containing only the V_1_ region of the complex improved resolution to 3.8-4.1 Å for this region by excluding the flexible peripheral stalks and V_o_ (membrane integrated) domain (Figs. S3, S4), although the d-subunit density forming the terminus of the rotor shaft was still poorly resolved (Fig. S3). Attempting to improve the clarity of the V_o_ domain by repeating this focussed refinement strategy using a mask for the V_o_ domain failed, even when falling back to a recombined particle set, masking out the V_1_ domain, and re-centring during initial 3D classification. Similarly, an attempt to better resolve the peripheral stalks by focussed refinement failed to converge, although this is unsurprising given that the V_1_ and V_o_ domains present a much more intense signal for alignment than the weak signal of the stalks.

### Examination of six states of EhV-ATPase at F-subunit position

The major three states of EhV-ATPase isolated were reconstructed to 4.2-4.4 Å (global) resolution (Figs. 2, S1, S3, S4) with the final maps consisting of between ~100,000-260,000 particles (15~37% of all particles) (Table 2). These three states correspond to the rotor orientations of conformations 1, 2 and 3 of TtV/A-ATPase (Nakanishi et al., 2018), and thus were defined as the major states 1, 2 and 3 with the orientations of the F-subunit (red asterisks in Fig. 2). These three major states were expected to rotate the rotor in a 120° rotation step to correspond to the pseudo three-fold symmetric rotary enzyme, but the cumulative rotation angle of 230° in State 3 was 10° smaller than the expected cumulative rotation of 240° (Fig. 2, Table 3).

**Figure 2.**
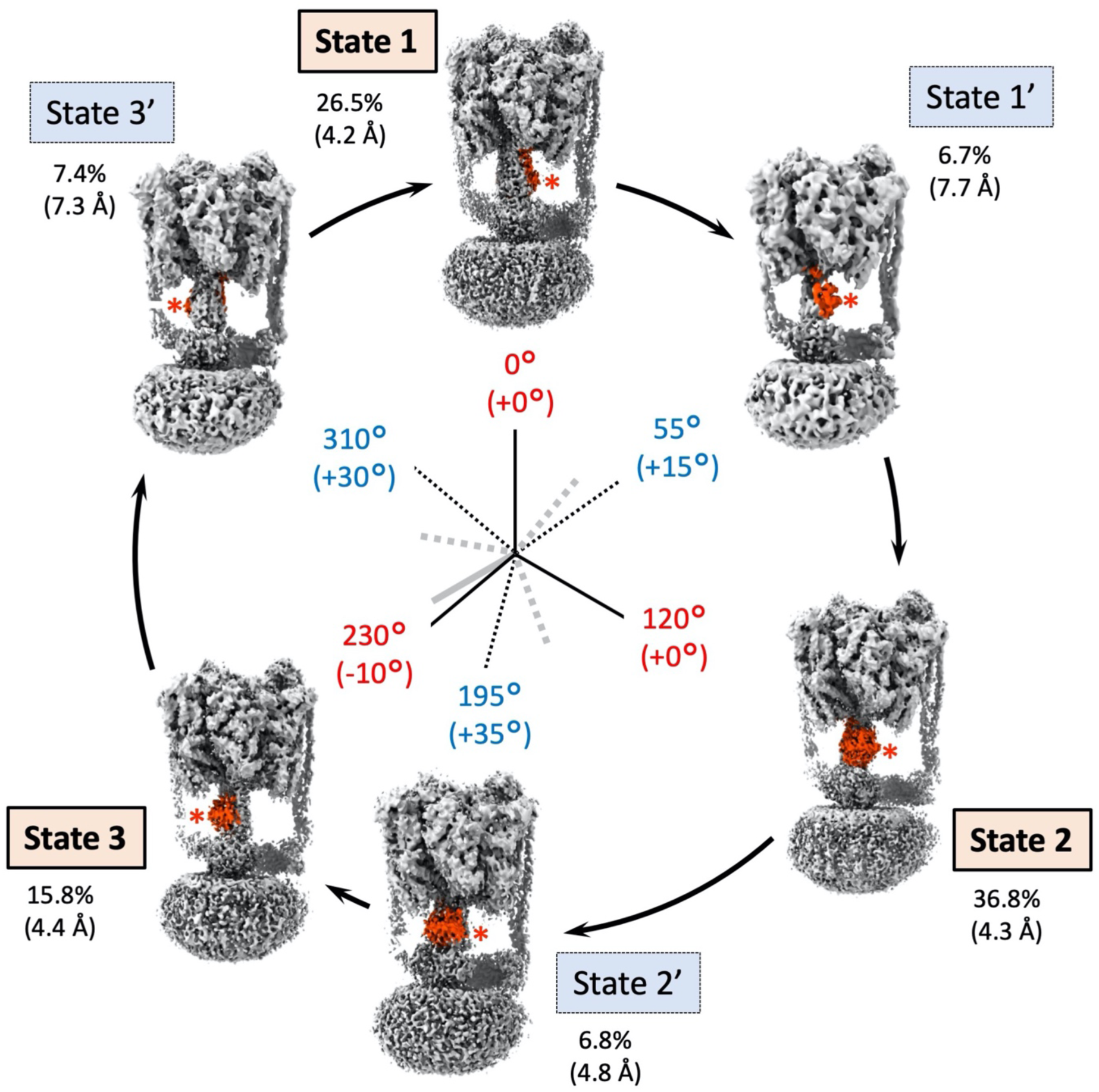
The six state structures of EhV-ATPase isolated in this study. The F-subunit position is highlighted in orange for easier identification of orientation of the rotor. Starting at State 1 at “12 o’clock” on the circle and proceeding clockwise as turnover proceeds when viewed from top to bottom. The six state structures are defined as State 1, State 1’, State 2, State 2’, State 3, and State 3’ with comparisons to the other V-ATPases. Total rotation of the rotor at F subunit is labelled in red for the major states and blue for the intermediate states internally of the circle. The gaps from the orientations based on the single molecular imaging studies (120° major pauses, and 40/80° subpauses in the major pauses) are in brackets. Relative percentages of the total final particles used and their respective resolutions (brackets) for each reconstruction are indicated externally of the circle. The cryo-EM maps of EhV-ATPase are located according to the orientation of the F-subunit.

**Table 2.**
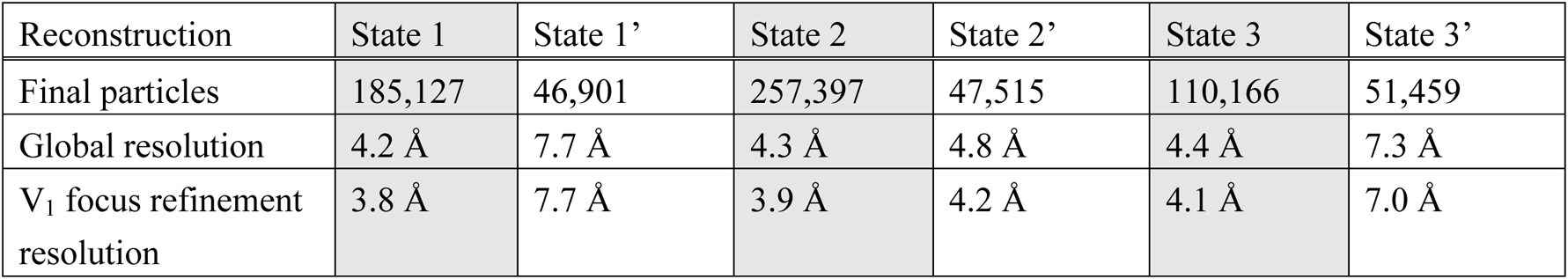
Statistics for the six 3D reconstructions.

| Reconstruction | State 1 | State 1' | State 2 | State 2' | State 3 | State 3' |
| --- | --- | --- | --- | --- | --- | --- |
| Final particles | 185,127 | 46,901 | 257,397 | 47,515 | 110,166 | 51,459 |
| Global resolution | 4.2 Å | 7.7 Å | 4.3 Å | 4.8 Å | 4.4 Å | 7.3 Å |
| $V_1$ focus refinement resolution | 3.8 Å | 7.7 Å | 3.9 Å | 4.2 Å | 4.1 Å | 7.0 Å |

**Table 3.** Orientations of the subunit at different positions in the complex during turnover, distances between the centres of d-subunit and c-ring, presence (or lack) of density in each A/B dimer ATP binding pocket, and distances from R573 in a-subunit to E139a or E139b in c-ring and their gaps.

|  | State 1 | State 1' | State 2 | State 2' | State 3 | State 3' |
| --- | --- | --- | --- | --- | --- | --- |
| Orientation angle of the D-subunit in V<sub>1</sub> domain | 0°<br>(0°) | 55°<br>(+15°) | 120°<br>(0°) | 195°<br>(+35°) | 230°<br>(-10°) | 310°<br>(+30°) |
| Orientation angle of the F-subunit in rotor | 0°<br>(0°) | 55°<br>(+15°) | 120°<br>(0°) | 195°<br>(+35°) | 230°<br>(-10°) | 310°<br>(+30°) |
| Orientation angle of the d-subunit in V<sub>o</sub> domain | 0°<br>(0°) | 55°<br>(+15°) | 120°<br>(0°) | 195°<br>(+35°) | 230°<br>(-10°) | 310°<br>(+30°) |
| Distance between the centres of c-ring and d-subunit at a slice | 3.7 Å | 3.9 Å | 7.6 Å | 9.4 Å | 3.1 Å | 2.0 Å |
| ATP pocket 1 | Empty | Full | Full | Full | Full | Full |
| ATP pocket 2 | Full | Full | Full | Empty | Empty | Full |
| ATP pocket 3 | Full | Full | Empty | Full | Full | Full |
| (a) E139a to R573 distance (Cα to Cα) | 11.9 Å | 11.4 Å | 12.3 Å | 11.0 Å | 11.8 Å | 10.6 Å |
| (b) E139b to R573 distance (Cα to Cα) | 9.6 Å | 10.5 Å | 9.6 Å | 10.8 Å | 9.8 Å | 11.6 Å |
| Gap (a – b) | +2.3 Å | +0.9 Å | +2.7 Å | +0.2 Å | +2 Å | -1.0 Å |

The three intermediate states named State 1’, State 2’, and State 3’, were identified by the orientation of the rotor with the F-subunit between the three major states of the EhV-ATPase (red asterisks in Fig. 2). These intermediate states are each composed of a similarly small number of particles (~50,000, Table 2). These states may be considered transitory because of the lower particle count (<7.4%) and the lower resolution (>4.8 Å) compared to the major three states (Figs. 2, S3, S4). Recent single molecular imaging studies of V_1_ domain indicated that there are subpauses divided by an approximately 40°/80° split within the major 120° rotation pauses of the pseudo three-fold symmetric rotation enzyme (Iida et al., 2019). The orientations of the rotor in the major states were close to these values in the 120° step, while the orientations of the rotor in the intermediate states were diverse (Fig. 2, Table 3). Compared to the expected +40° subpause from the previous major step, the 55°, 195°, and 310° rotor orientations at the F-subunit of States 1’, 2’, and 3’ showed subpause shifts of +15°, +35°, and +30°, respectively (Table 3).

### Examination of six states of EhV-ATPase at V_1_ position

The orientations of the rotor D-subunit based upon the direction of the two long helices showed the same angles as the position of the rotor F-subunit in both the major and intermediate states in the V_1_ domain (red boxes in Fig. 3, Table 3). The result shows the rotor D-subunit is not elastic but rigid between the V_1_ domain and the F-subunit. The cryo-EM maps at the resolution from 7.3 to 4.2 Å clearly illustrated the conformations of the A/B subunit in V_1_ domain in the pseudo three-fold symmetric rotary enzyme. These are easily classified as “Open”, “Closed”, and “Semi-closed” based on the previous report of the crystallographic models of the V_1_ domain (Suzuki et al., 2016), each showing the different ATPase catalytic condition. In the major three states, each A/B subunit takes a different conformation, which is switched to rotate in the order of “Open”, “Semi-closed”, and “Closed”, in a continuous cycle between the three states, rotating the central rotor (Fig. 3). Interestingly, in the intermediate V_1_ states, State 1’ and State 3’ took the similar conformation with State 1 and State 3, respectively, while State 2’ showed a similar conformation to State 3 (Fig. 3).

**Figure 3.**
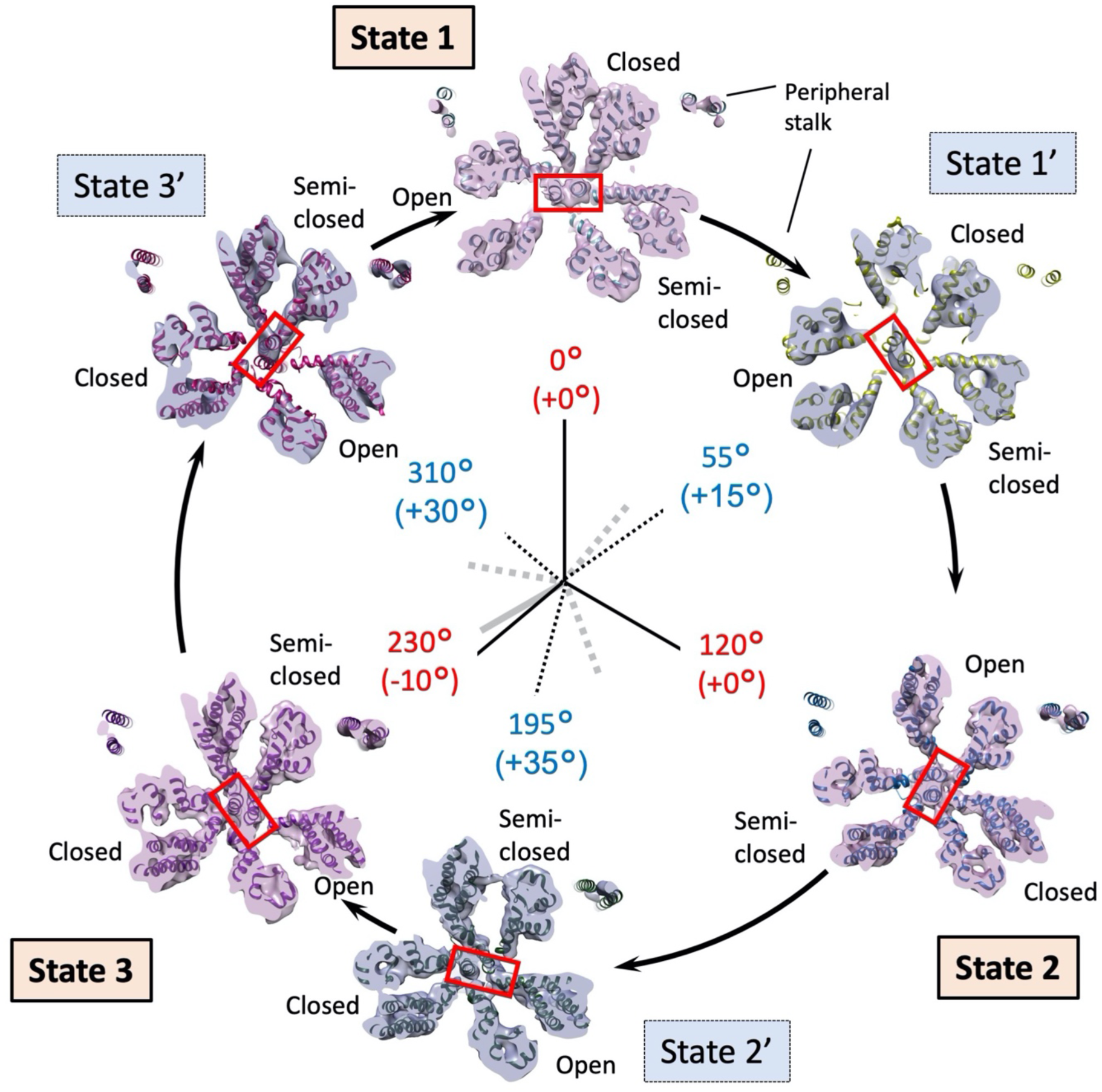
V_1_ domain cross-section view of the six states. The figure is laid out as shown in Fig. 2 viewed from the V_1_ to V_o_ domains. The rotor (D subunit) is boxed in red, demonstrating the positions of a pair of the longest helices. The conformations of the A/B subunit in V_1_ domain are indicated with “Open”, “Closed”, and “Semi-closed”. The positions of peripheral stalk are labelled.

### Examination of six states of EhV-ATPase at V_o_ position and their off-axis assembly

The orientation angles of the rotor shaft terminus in the six states of EhV-ATPase were estimated with the d-subunit located between D-subunit and c-ring (Fig. 1), which also showed angles similar to the rotor F-subunit position at V_o_ position in both major and intermediate states (Fig. 4, Table 3). The result indicated the rotor D-subunit and the following rotor shaft terminus d-subunit are not elastic but rigid between the V_1_ domain and the V_o_ domain.

**Figure 4.**
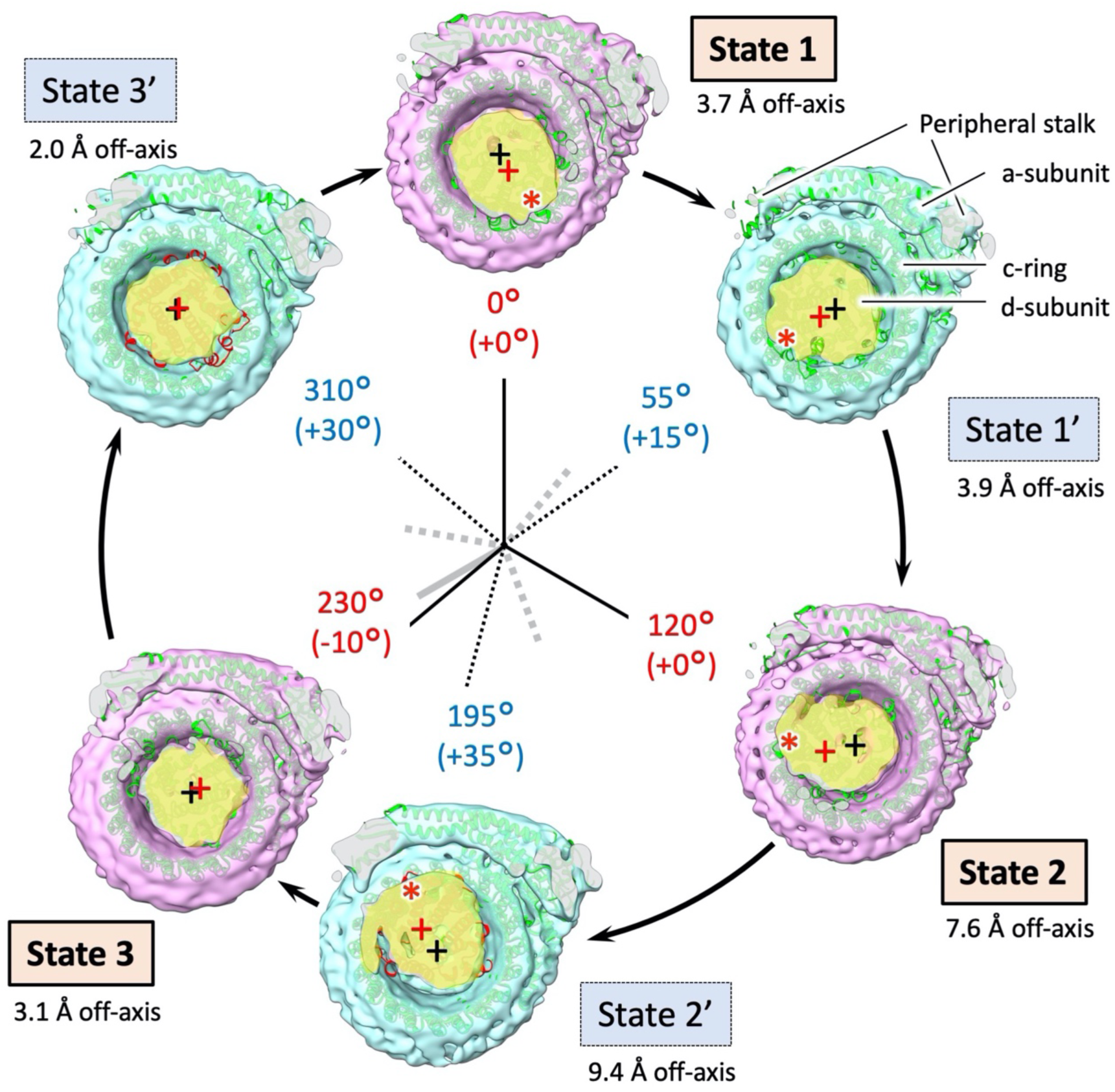
V_o_ domain cross-section view of the six states. The figure is laid out as in Fig. 2, viewed from V_1_ to V_o_ domains. The cross-sections of the d-subunit are displayed in yellow. The centre of the c-ring on each state is marked with a black cross, while the centre of the d-subunit cross-section is marked with a red cross. Deviation of the d-subunit from the centre of the c-ring is indicated by Å.

On the other hand, the off-axis nature of the rotor varied during turnover of the EhV-ATPase complex (Fig. 4). The degree to which the rotor is “off-axis” changed continuously as rotation proceeds. For State 2, and State 2’, the off-axis nature of the rotor was clearly detected, while for State 3 and State 3’ the centre of the d-subunit is fairly close to the centre of the c-ring (red and black crosses in Fig. 4, Table 3). Slicing these maps horizontally at a V_o_ domain position and measuring the distance from the centre of the c-ring to the centre of the d-subunit, the larger off-axis centres of d-subunit were detected as 7.6 and 9.4 Å in State 2 and State 2’, while the smaller off-axis centres of the d-subunit were detected as 3.1 Å in State 3, and 2.0 Å in State 3’. The intermediate off-axis centres of d-subunit were identified as 3.7 Å in State 1 and 3.9 Å in State 1’ (Fig. 4, Table 3). This off-axis centring of the d-subunit can be caused by the interaction between the large c-ring and the small d-subunit (red asterisks in Fig. 4), although the interaction residues between these subunits were not cleared at the limited resolution of the V_o_ domain. This can be further partially interfered by the extrinsic domain of the a-subunit connected to the peripheral stalks to support the V_1_ domain (Fig. 5). Interference between the d-subunit of the rotor and the extrinsic domain of the a-subunit were observed in State 2’ and State 3 (blue asterisks in Fig. 5), where the large off-axis nature of the d-subunit (9.4 Å off-centre) in State 2’ was significantly centred (3.1 Å off-centre) in State 3 (Fig. 4).

**Fig. 5.**
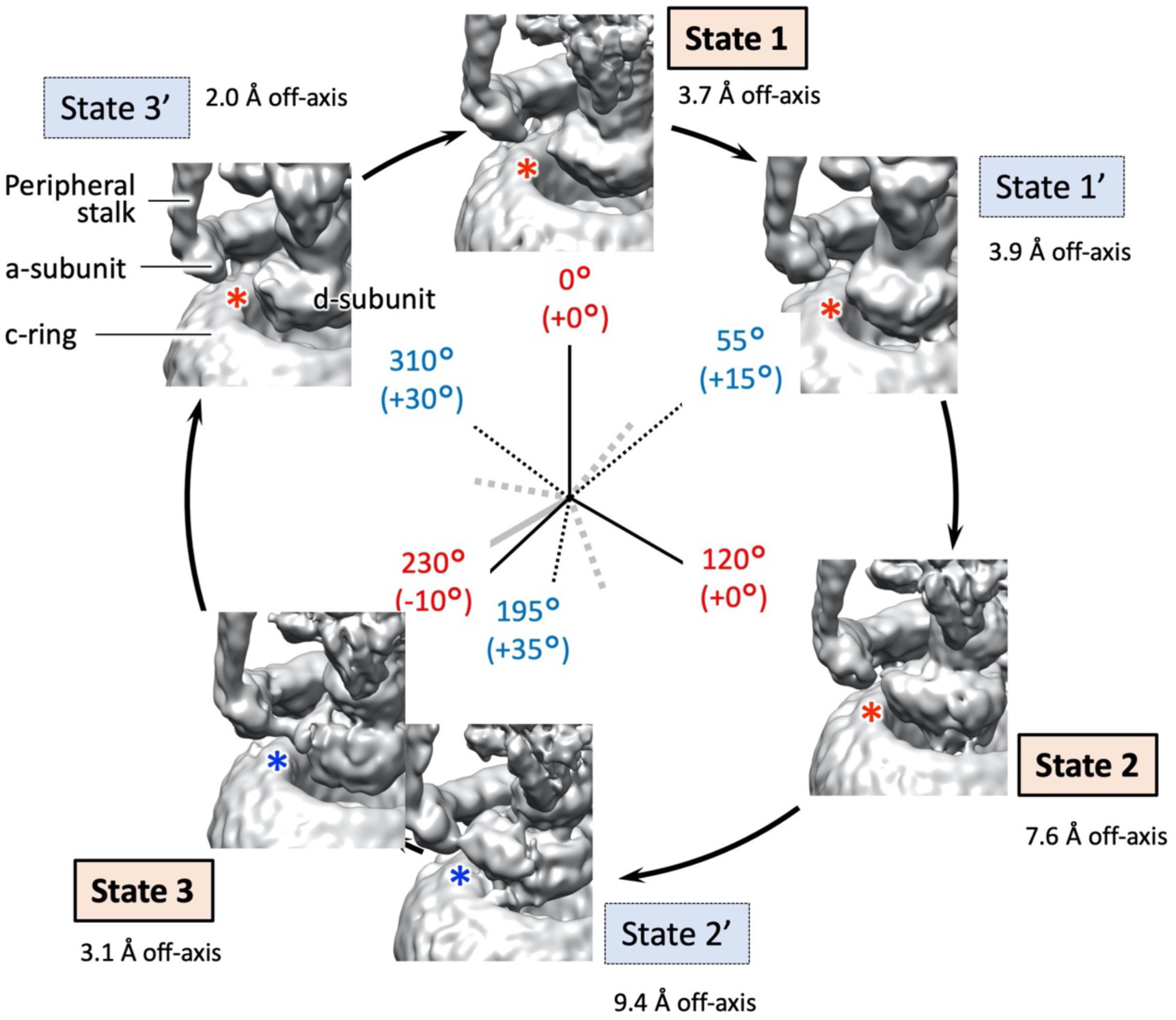
Isosurface view of the reconstructions for each state focussed on the rotor and terminus of the a-subunit arm. The figure is laid out as in Fig. 2, viewed down from V_1_ to V_o_ domains. The maps were filtered by their local resolution. A red asterisk indicates no interaction (density bridge) between the rotor and the a-subunit arm, a blue asterisk indicates an interaction (density bridge). Rotational state of rotor indicated for each state. Total rotation of the rotor at F subunit, with the change from previous orientation in brackets, is labelled in red for the major states and blue for the intermediates. The off-axis distances shown in Fig. 4 are indicated externally of the circle.

### Examination of six states of EhV-ATPase in the ATP binding pocket

Significant studies have been focussed on the turnover of the V_1_ domain for EhV-ATPase, permitting the development of models of how turnover proceeds mechanistically (Iida et al., 2019; Iino et al., 2015; Ueno et al., 2014) and structurally (Arai et al., 2013; Suzuki et al., 2016) during ATP hydrolysis. In this study, a cross-section of the maps across the three nucleotide binding pockets in the V_1_ domain directly show that in the major three states, two nucleotide binding pockets contain densities for nucleotides (Fig. 6, blue circles in States 1, 2, and 3), but the third one is empty (Fig. 6, red circles in States 1, 2, and 3). In Fig. 6, the crystallographic model of PDBID: 5KNC was fitted into the maps to see whether density for a bound nucleotide is present or not. These images are magnified in Figs. S5, S7, and S9, respectively. In addition, these three states exhibited continuous changes in binding/catalytic dwell during turnover (States 1, 2, and 3 in Fig. 6). On the other hand, in the intermediate States 1’ and 3’, all three nucleotide binding pockets contains density (Fig. 6, blue circles in State 1’ and 3’), while State 2’ shows the similar binding coordination as State 3 (State 2’ in Fig. 6). These images are magnified in Figs, S6, S8, and S10, respectively. These results are consistent with the notion that the major three states correspond to the main pauses waiting for ATP binding, observed in the previous single molecular study of isolated V_1_ domain (Iida et al., 2019).

**Fig. 6.**
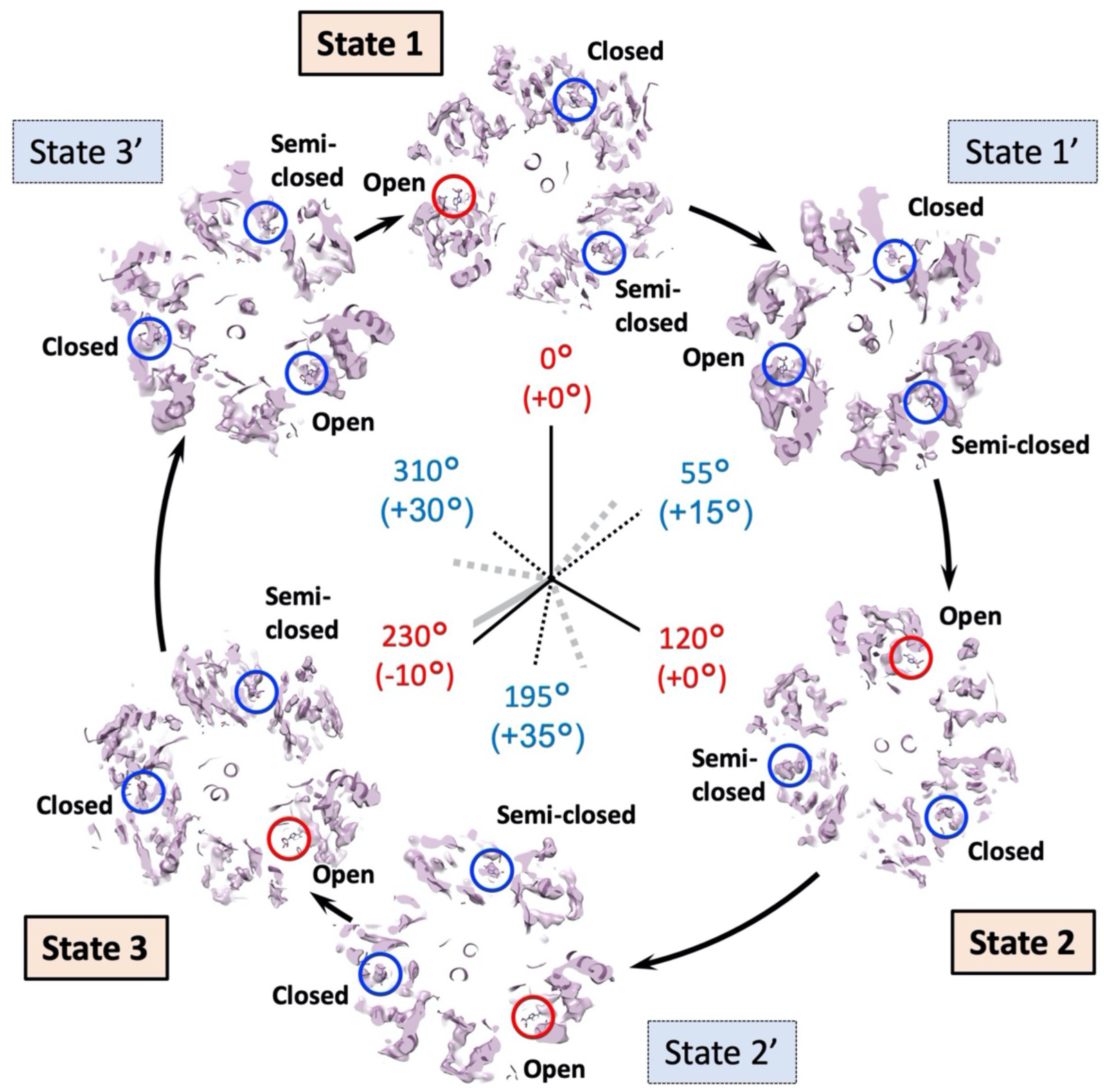
The ATP binding pocket densities of the different states in EhV-ATPase. The figure is laid out as in Fig. 3, viewed down from V_1_ to V_o_ domains. The maps are sliced through the density map at the level of the ATP binding pocket, and with PDBID:5KNC fitted to check whether density for a bound nucleotide is present or not. Blue circles indicate the presence of density corresponding to the bound nucleotide in the fitted PDB, red circles indicate missing density (only nucleotide from PDB is visible). Rotational state of rotor indicated for each state of six. Cumulative rotation angles of the rotor at the V_1_ domain are labelled as the same as Fig. 3.

## Discussion

Here we identified six independent conformations of EhV-ATPase by cryo-EM using an active sample which was encouraged to turn over through the addition of ATP, Mg^2+^ and Na^+^. While the purified EhV-ATPase complex is fragile, and the actively rotating complex even more so, the identification of three major states and three intermediate states provides some further structural clues to the activity of this so far unique “off-axis” adaptation to a ubiquitous membrane protein complex.

Single molecular imaging studies of the isolated V_1_ domain of EhV-ATPase have indicated that there are three major pauses of each 120° rotation of the rotor of the pseudo three-fold symmetric rotary enzyme and three intermediate subpauses, each divided by approximately 40°/80° split within the major pauses (Iida et al., 2019). We directly examined these rotor angles of the major three states and the novel intermediate three states in the six structures. The results show the major pauses for each 120° rotation in State 1 and State 2, but the cumulative angle of 230° in State 3 was 10° smaller than the expected cumulative angle of 240° at all rotor positions (Figs. 2, 3, and 4). This may be caused by the interference between the off-axis rotor and the stator subunit in the EhV-ATPase. The rotor shaft terminus of the d-subunit and the c-ring in the V_o_ domain showed relatively lower resolution, but when applying local-resolution filtering a density bridge was identified between the d-subunit and the extrinsic arm of the a-subunit (blue asterisk in State 3 in Fig. 5). The molecular interaction between these subunits may cause interference in the main pause of State 3. As previous work utilised only V_1_ domain (Suzuki et al., 2016), steric hindrance from the membrane-extrinsic region of the a-subunit would not be observed. Furthermore, the same differential in this direction of the rotor in State 3 were found at all positions in the V_1_ domain, rotor, and V_o_ domain (Table 3). This suggests that the rotor coupling between V_o_ and V_1_ is not elastic but rigid as reported by Otomo et al. using single molecular imaging studies (Otomo et al., 2022).

On the other hand, the orientational gap of the subpauses was more diverse in the intermediate states. The smallest gap from the expected subpause angle of 40/80° in the 120° rotation was shown in State 1’, where the subpause angle of 55° showed a positive gap of 15° at all rotor positions (Table 3). In contrast, the subpause angles of 195° and 310° in State 2’ and State 3’ showed positive gaps of 35° and 30°, respectively, and were significantly larger than that of State 1’ (Table 3). Eventually, the subpauses were caused at 55°/65° in State 1’, 75°/35° in State 2’, and 80°/50° in State 3’, respectively, in the whole EhV-ATPase complex (Table 3). Compared to the subpause angle observed in the isolated V_1_ domain (Iida et al., 2019), it was suggested that the intramolecular interactions in the whole complex could produce significantly different angles in each state. Furthermore, in State 2’, the subpause angle (+35° from the expected subpause angle) may reflect the interference of the density bridge between d-subunit and the extrinsic arm of the a-subunit (blue asterisk in State 3 in Fig. 5), similar to the major pause gap of −10° in State 3 mentioned in the previous paragraph (Table 3).

The off-axis rotor assembly was observed between the d-subunit and the c-ring (Fig. 4). We measured the distance between the center of c-ring and the center of the d-subunit in the horizontal cross-section of the V_o_ domain (Fig. 4, Table 3). The smallest gap distance of 2.0 Å observed in State 3’ was gradually increased during turnover, and finally the largest gap distance of 9.4 Å appeared in State 2’. After that, the gap distance gradually decreased again toward the smallest 2.0 Å in State 3’. When the off-axis motion was compared with the density bridge between the rotor shaft terminus d-subunit and the extrinsic domain of the a-subunit we observed (Fig. 5), the interreference can be related to the off-axis rotor motion. The gap distance with the off-axis motion increases when the interaction between the d-subunit and the c-ring moves to the opposite side of the a-subunit (Figs. 4 and 5, red asterisks in State 1 to State 2). However, the off-axis distance decreases after the interference between the d-subunit and the extrinsic arm of the a-subunit (Fig. 5, Table 3). Finally, the off-axis motion was minimized at State 3’ (Fig. 4). The off-axis motion initially caused by the interaction between the d-subunit and the c-ring is likely to be re-centered with the interference between the d-subunit and the extrinsic arm of the a-subunit.

State 2’ is unique among the intermediate states and has only a small (global) resolution loss compared to the major states, while State 1’ and State 3’ are markedly lower resolution (>7 Å) (Figs. S3, S4, Table 2). Despite containing a comparable number of particles to States 1’ and State 3’, State 2’ is still <5 Å. In State 2’, the d-subunit forming the rotor shaft terminus is approaching the extrinsic arm of the a-subunit (Fig. 5), furthest from the transition of the a-subunit from membrane-intrinsic to membrane-extrinsic. This region is, frustratingly, the weakest point of many of the reconstructed maps, indicating that it is highly mobile, but the State 2’ reconstruction shows a potential interaction between the d-subunit and the a-subunit at this position when the map is filtered by local resolution (blue asterisk in State 2’ in Fig. 5). This may contribute to the improved stability in addition to the re-centring of the rotor, and corresponding improvement in resolution of State 2’ compared to State 1’ or State 3’ despite the relatively low particle count.

Based on these results and observations of the ATP binding pocket in V_1_ domain (Fig. 6, Figs. S5-S10) and those of the previous structural and single molecular studies of isolated V_1_ domain (Iida et al., 2019; Suzuki et al., 2016), we have built a catalytic model of EhV-ATPase through six states (Fig. 7). The process involved in a single rotation of the rotor within the V_1_ domain consists of three repeats between three major states, each of which includes a phosphate release, an ATP binding, an ADP release, and an ATP hydrolysis events. These reactions would correspond to a conformational shift in the associated A/B dimer, and in turn would affect the rotation of the rotor. For example, looking at the event that occurred between State 1 and State 2, there is a clear distinction between the processes from State 1 to State 1’ and State 1’ to State 2. In the early process from State 1 to State 1’, inorganic phosphate is released from the hydrolyzed ATP (ADP + Pi) in the closed pocket, and new ATP binds to the open pocket. During this period, the structure of the pockets of “open”, “closed”, and “semi-closed” remains similar, and the rotor rotates 55°. Then, in the later process between State 1’ and State 2, the ADP is released from the closed pocket and the structure of the pocket changes from “closed” to “open”. The “semi-closed” pocket changes to “closed”, and ATP is hydrolyzed to ADP and Pi. The open pocket containing ATP turns into “semi-closed”. During this period, the rotor rotates 65°. A similar model can be applied to the process between State 3 and State 1, but the rotation angle of the rotor is different: 80° between State 3 and State 3’ and 50° between State 3’ and State 1. Differences in the rotor angle of the rotation can be caused by the rotor’s off-axis assembly, which is clearly observed at the V_o_ domain level (Fig. 4).

**Figure 7.**
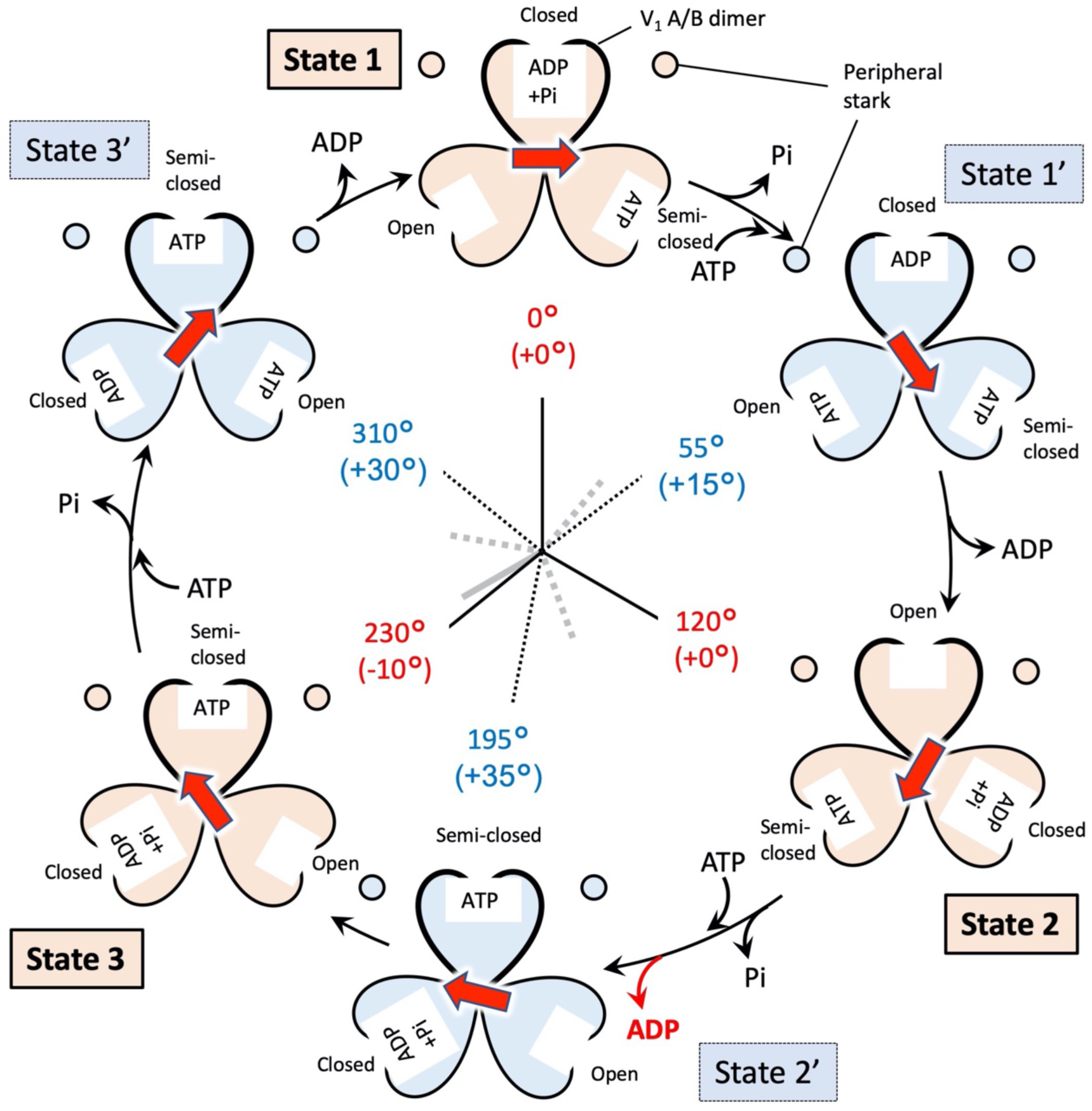
Schematic overview of the turnover of EhV-ATPase. The V_1_ domain is shown by the six states of the major and intermediate conditions, when viewed from V_1_ to V_o_ domains. Rotor orientation is indicated with a red arrow. In each lobe of the V_1_ trimer is indicated whether the nucleotide binding pocket is empty, or containing ATP, ADP or ADP+Pi. The figure is laid out as in Fig. 6.

The special case of the model was observed between State 2 and State 3 (Fig. 7). The structure of the ATP binding pockets and the nucleotide densities of State 2’ are similar to State 3 of the major state except for the rotor angle (Figs. S8 and S9). The rotor angle of rotation in State 2’ indicates that it is in the intermediate state, in which the 120° major rotation of the rotor was divided into 75°/35°. This structure indicates that the all catalytic events for Pi release, ATP binding, ADP release, and ATP hydrolysis occur in the early process between States 2 and State 2’, and only rotor rotation occurs in the later process between State 2’ and State 3. If that is true, why does the rotor rotation occur between State 2’ and State 3, in which period, no chemical events occurs? The abnormal behaviour can be caused by the interference between the d-subunit and the extrinsic arm of the a-subunit described above (Fig. 5). The interactions between these subunits that cause re-centring of the off-axis rotor between State 2’ and State 3 may cause rotor rotation without chemical events. Unfortunately, while the presence or lack of nucleotide in a binding pocket is easy to determine, differentiating between ATP, ADP with inorganic phosphate and ADP without inorganic phosphate is considerably more difficult and will require extremely high resolutions in the V_1_ domain. This mechanism is similar to that recently proposed by Shekhar et al. (Shekhar et al., 2021). State 2’ not only exhibits a binding state closer to that of the binding/catalytic dwell than the ADP release dwell, but the rotor has more advanced significantly than State 1’ with a 75° rotation. State 3’ shows a similarly advanced rotation of 80° when compared to State 1’ of the 55° rotation, but presents the densities corresponding to three possible bound nucleotides. This indicates one of the densities is in the ADP release dwell, and is thus consistent with the previously proposed model of turnover (Suzuki et al., 2016).

When the EhV-ATPase is activated, the relative position between the a-subunit and the c-ring changes continuously within the membrane (Fig. 8, Table 3). Despite our best efforts, we have thus far been unable to improve the clarity of the V_o_ domain beyond that exhibited in the whole-complex reconstructions. We attribute this to the use of Na^+^, Mg^2+^ and ATP to generate turnover prior to vitrification, causing the c-ring to be present in multiple rotational states within the detergent, even within the distinct classes. It is possible to fit the α-helices to the density (which comprises most of the c-ring and a-subunit) but clarifying side-chain identity and orientation is essentially impossible. Therefore, when examining the V_o_ domain, we measured the distance between the c-ring and the a-subunit from the Cα of the residues of interest: R573 and R629 for a-subunit, E139 for c-ring, where the residue of E139 is suggested to interacts with vertically aligned two residues of R573 and R629 in a-subunit and prevent the ion shortcut without c-ring rotation (Murata et al., 2005; Otomo et al., 2022; Zhou and Sazanov, 2019). However, in the current resolution in V_o_ domain, two possibilities for the E139 location must be considered in the c-ring, because the residue of E139 is positioned in one of two transmembrane helices located the outer c-ring. Therefore, we firstly measure the distances between the residue R573 of the a-subunit and two possible resides of E139 locations temporally labelled as E139a and E139b in Fig. 8. As a result, the major three states show the average distance of 9.7±0.1 Å from the residue R573 of the a-subunit to the residue E139a, and the average distance of 12±0.3 Å to the residue E139b (Fig. 8, Table 3). These distances are approximately consistent in all three major states, where the one of two helices locates closer to the residue R573 of the a-subunit. The intermediate three states, on the other hand, shows that both helices are at more similar averaged distances: 11±0.6 Å for the helix containing residue E139a and 11±0.4 Å for the helix containing residue E139b. For the residue R629, we also had a similar result. Previous reports showed that each monomeric c-ring member binds a single Na^+^ ion in a pocket consisting of the coordinating residues L61, T64, Q65, Y68, Q110 and E139 (Murata et al., 2005), where Y68 coordinates with E139 rather than directly with bound Na^+^. Further, E139 plays a significant role in the binding of Na^+^ for translocation from one side to the other of the membrane (Kawano-Kawada et al., 2012) with mutagenic experiments to E139(D/Q) eliminating Na^+^ transport, while E139D still permitted Li^+^ transport (Kawano-Kawada et al., 2011). On the other hand, R573 of the a-subunit is proposed to act as a “latch” for c-ring rotation (to prevent c-ring rotation in the wrong direction, and thus a shortcut of the Na^+^ ion transport) (Mitome et al., 2010). The equivalent residue of the TtV/A-ATPase is R563 of the a-subunit, and was proposed to act to prevent this shortcut in *T. thermophilus* (Zhou and Sazanov, 2019). These observations suggest that in the major state both binding and release of transported Na^+^ positively occur between R573 of the a-subunit and E139 in the c-ring. By contrast, in the three intermediate state the c-ring would be in an intermediate stage of rotation, with no binding pocket directly accessible to either the ion ingress or egress pores.

**Figure 8.**
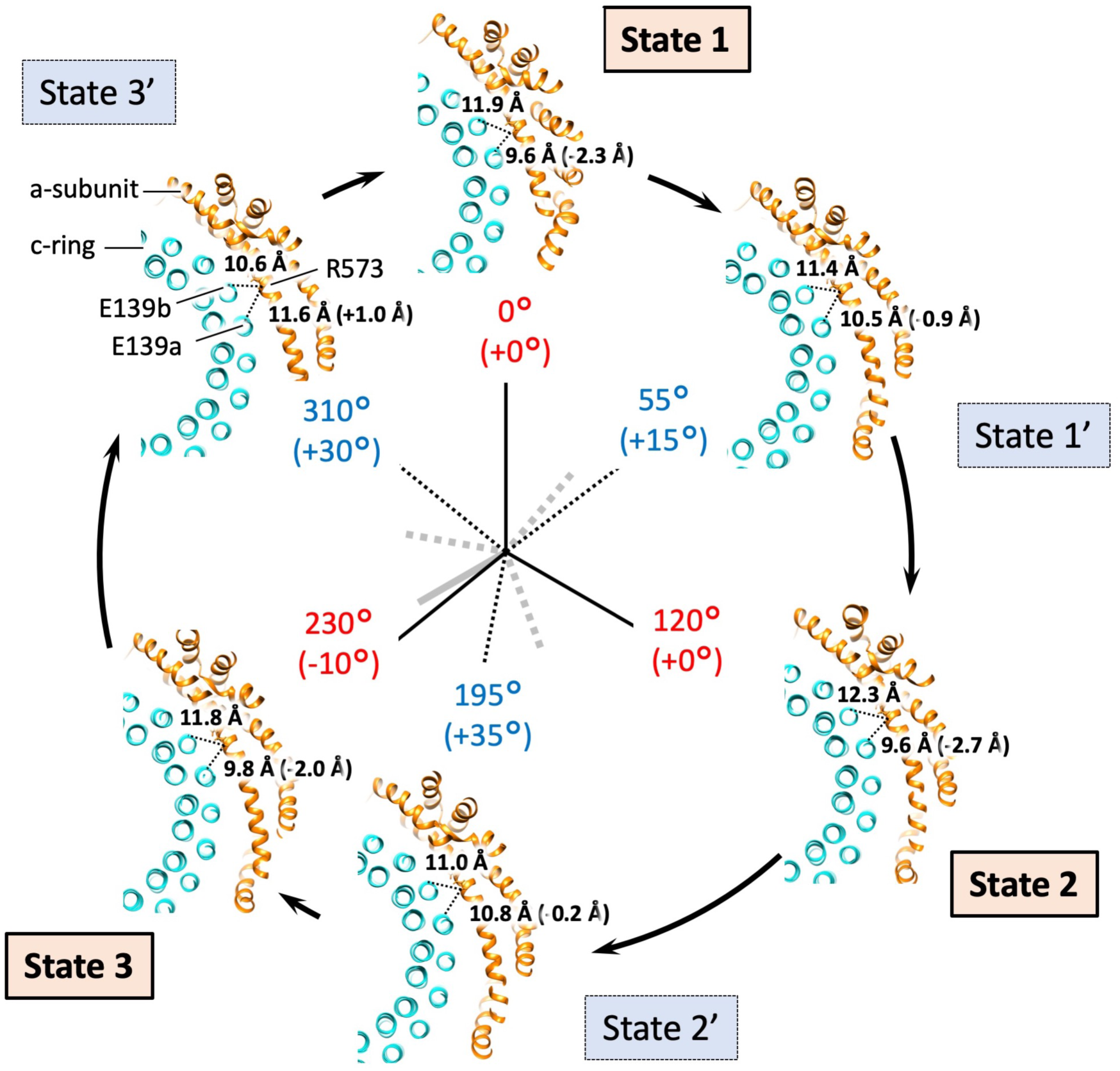
Focussed view of the R573/E139a or E139b locale of the six states. The figure is laid out as in Fig. 2, viewed down from V_1_ to V_o_ domains. Due to resolution constraints (Figs. S3 and S4) it is not possible to reliably determine residue side-chain positions, thus distances are measured (shown with a dashed line) from Cα of E139a and E139b of the c-ring to Cα of R573 of the a-subunit. E139a and E139b are one of two possible E139 positions on the two outer transmembrane helices, respectively. Absolute distance for each state is indicated, with difference between the two in black brackets. Also see Table 3.

## Conclusion

In this study, we report the structures of the six states of EhV-ATPase, which consisted of three major states and three intermediate states. EhV-ATPase showed a variety of rotor orientations in one major state and three intermediate states. These were suggested to be caused by the off-axis rotor assembly and its interreference with the extrinsic arm of the a-subunit. Densities in the nucleotide binding pockets in the motor V_1_ domain showed the different chemical states between the major and intermediate states. Especially, State 2’ state show a novel state, in which the chemical state is similar to the next State 3, but the rotor orientation shows that of subpause. The large c-ring of EhV-ATPase has advance to accurately transport ions with dual helices per ion, compared to the small c-ring with single helix per ion of the TtV/A-ATPase. However, in addition to the asymmetric dual peripheral stalks, the mismatch between the relatively smaller d-subunit and the larger c-ring likely causes the unique off-axis assembly of the rotor complex in EhV-ATPase. The structures of the eukaryotic V-ATPases revealed that additional subunits in the c-ring and the symmetric triplet peripheral stalks perform the on-axis rotor assembly even in the large c-ring (Marshansky et al., 2014). EhV-ATPase may show the evolutionary pathway of the ubiquitous membrane protein complex family of V-ATPases from prokaryotes to eukaryotes. Further investigations are necessary to elucidate this relationship between V-ATPases.

## Experimental Procedures

### Expression and purification of recombinant EhV-ATPase

Expression and purification of recombinant EhV-ATPase were performed as previously described (Tsunoda et al., 2018). In brief, *E. hirae* V-ATPase was expressed recombinantly in *Escherichia coli* and purified by affinity purification with a Ni^+^-NTA (nitrilotriacetic acid) column (Ni^+^-NTA Superflow; Qiagen, Hilden, Germany). The column was preconditioned with buffer consisting of 50 mM potassium phosphate, 100 mM KCl, 5 mM MgCl_2_, 20 mM imidazole, 10% glycerol, 0.05% n-dodecyl-β-D-maltoside (βDDM) at pH 7.5. The column was washed, and protein was eluted with buffer containing 50 mM potassium phosphate, 100 mM KCl, 5 mM MgCl_2_, 300 mM imidazole, 10% glycerol, 0.05% βDDM at pH 7.5. The purified protein was concentrated using an Amicon Ultra 100 K filter (Merck Millipore, Billerica, Massachusetts, USA) before further purification using Superdex 200 gel filtration (GE Healthcare, Little Chalfont, UK) preconditioned with 50 mM Tris-HCl, 5 mM MgCl_2_, 10% glycerol, 0.05% βDDM at pH 7.5.

### Cryo-EM grid preparation

Purified EhV-ATPase was mixed with ATP and Na^+^ to final concentrations of 0.13 mM (protein), 7 mM (ATP) and 100 mM (Na^+^) before vitrification by plunge freezing using an Vitrobot Mark IV (Thermo Fisher Scientific, Hillsboro, Oregon, USA) onto Quantifoil R1.2/1.3 copper grids (Quantifoil Micro Tools GmbH, Großlöbichau, Germany) which had been glow discharged immediately beforehand.

### Cryo-EM data acquisition

Micrograph movies were acquired on a JEOL CRYOARM 300 microscope (JEOL Ltd., Tokyo, Japan) equipped with a Gatan K3 direct electron detector (Gatan Inc., USA) at a sampling scale of 1.01 Å/pixel using SerialEM automation (Mastronarde, 2005). 50 frames were recorded for each movie at low dose conditions for a total dose of 50 e^−^/Å^2^. Three good grids were used for data acquisition. Micrographs were collected in three rounds; 19,250 micrographs in the first session and a further 15,978 micrographs collected in a second round. Due to strong orientation preference, a further round of data collection was carried out at 20° stage tilt and 30° stage tilt collecting a further 5,032 and 3,112 micrograph movies at those respective tilts. As the Gatan K3 suffered from significant gain drift during data acquisition, gain references were generated by summing raw movie frames using the cisTEM (Grant et al., 2018) program ‘sum_all_tif_files’ in appropriate batches required to avoid poor gain correction. Movies were motion corrected using MotionCor2 (1.3.1) (Zheng et al., 2017) and the contrast transfer function (CTF) estimated using CTFFIND (4.1.10) (Rohou and Grigorieff, 2015). Data collection and image processing information are summarized in Table 1.

### Cryo-EM data analysis

A previously reported reconstruction (Tsunoda et al., 2018) was used to generate projections for a first-round automated particle picking on a subset of micrographs with Gautomatch (Zhang, 2016). These picked particles were extracted, 3× down-sampled and 2D classified. Clear classes were used for input into the RELION 3.1 (Scheres, 2012; Zivanov et al., 2020) autopicking routine. Due to difficulties visualising EhV-ATPase without phase plate (Tsunoda et al., 2018), generous settings were used for autopicking resulting in a total of 4,383,273 particle candidates picked. Particles were picked from micrographs grouped into ~1,000 randomised micrographs per group, particles extracted with 3× downsampling and 2D classified in batches. Each batch picked approximately 90-110,000 particles, and between 60-80,000 particles were carried forward per batch in “good” classes. The good classes from every five batches were grouped together and passed to 3D classification with an angular sampling of 7.5° into 15 classes, where classes which were evidently complete complexes were selected. Classes with a damaged V_o_ (membrane) domain, or broken rotor shaft were discarded. Particles were combined and 3D refined with a soft mask to improve angular assignments, before further 3D classification with alignment disabled into 25 classes. These classes were selected and grouped based on the general position of the rotor before a further 3D refinement with a reference generated by “relion_reconstruct” using each particle selection and a mask generated from that reference, which further optimised the angular assignment. Further 3D classification was carried out on each set into 25 classes, and each class manually assigned to one of six states based on rotor position (State 1, 1’, 2, 2’, 3 or 3’) (Fig. S1).

A soft mask focussed on the V_1_ domain and rotor shaft, excluding the peripheral stalks and V_o_ domain, was generated and a 3D refinement carried out once more. After this a 3D classification using the same mask was carried out into 25 classes with alignment off. This resulted in seven classes with estimated resolutions exceeding 8 Å, which were selected and classified once more into 25 classes. States 1 and 3, the two positions either side of State 3’ were combined, and 3D classified once more into 15 classes with classes grouped based on rotor position. These class assignments were passed to individual 3D refinement. Particles were re-extracted and re-centred at full scale, with “relion_reconstruct” used to generate references of them with the maximum resolution set to 15 Å. Reconstructions of the complete complex in each state were carried out, with CTF refinement, resulting in final resolutions of shown in Table 2 and Fig. S4. Localised masks were generated of the V_o_ and V_1_ domains, and further reconstructions carried out. This improved resolution for V_1_ domain, but not for V_o_ domain (Figs. S3 and S4).

### Visualisation

Micrographs, particles and 2D classes were visualised using the “relion_display” module of RELION 3.1 (Zivanov et al., 2020). 3D maps and PDB models were visualised using IMOD (Kremer et al., 1996), UCSF Chimera (Goddard et al., 2007; Pettersen et al., 2004) or UCSF ChimeraX (Goddard et al., 2018).

### Model building

The V_1_ domain crystal structures of *E. hirae* V-ATPase (PDBID: 3VR6) (Arai et al., 2013) and (PDBID: 5KNB, 5KNC) (Suzuki et al., 2016) were used for the V_1_ domain, and the crystal structure of the c-ring (PDBID: 2BL2) (Murata et al., 2005) was used as a component of V_o_. The a-, d-, E- and G-subunits were generated by homology modelling using the I-TASSER suite (Yang and Zhang, 2015) and fitted against independently generated cryo-EM maps (described above) of the varying states.

### Data availability

The cryo-EM maps of the entire EhV-ATPase complex have been deposited in the Electron Microscopy Data Bank under accession number EMD-#### for State 1, EMD-#### for State 1’, EMD-#### for State 2, EMD-#### for State 2’, EMD-#### for State 3, and EMD-#### for State 3’. The cryo-EM maps of the V_1_ domain of the EhV-ATPase complex have been deposited in the Electron Microscopy Data Bank under accession number EMD-#### for State 1, EMD-#### for State 1’, EMD-#### for State 2, EMD-#### for State 2’, EMD-#### for State 3, and EMD-#### for State 3’.

## Acknowledgements

We thank Fumiaki Makino at JEOL, and Takayuki Kato at Osaka University, for helping data collection by CRYO ARM 300 (JEOL Inc.), and Akihiro Otomo at Institute for Molecular Science for helpful discussion. This study was supported by Interdisciplinary research project by young researchers in NINS (to CS), JSPS KAKENHI Grant Number JP16H06280 (to RI), Platform Project for Supporting Drug Discovery and Life Science Research (Basis for Supporting Innovative Drug Discovery and Life Science Research (BINDS)) from AMED under Grant Number JP17am0101001 (to KM), and the Grant-in-Aid for Scientific Research on Innovative Areas “Molecular Engine” (JP19H05380 to HU, JP18H05425 to TM, and JP18H05424 to RI).

## Author Contributions

KM conducted and designed the experiments; HU, TM, and RI prepared the specimens. CS collected the cryo-EM data. RNBS process the cryo-EM images. RNBS and KM wrote the paper. All authors revised the paper.

