## Supplemental Figure 1-10 for "Six states of *Enterococcus hirae* V-type ATPase reveals non-uniform rotor rotation during turnover"

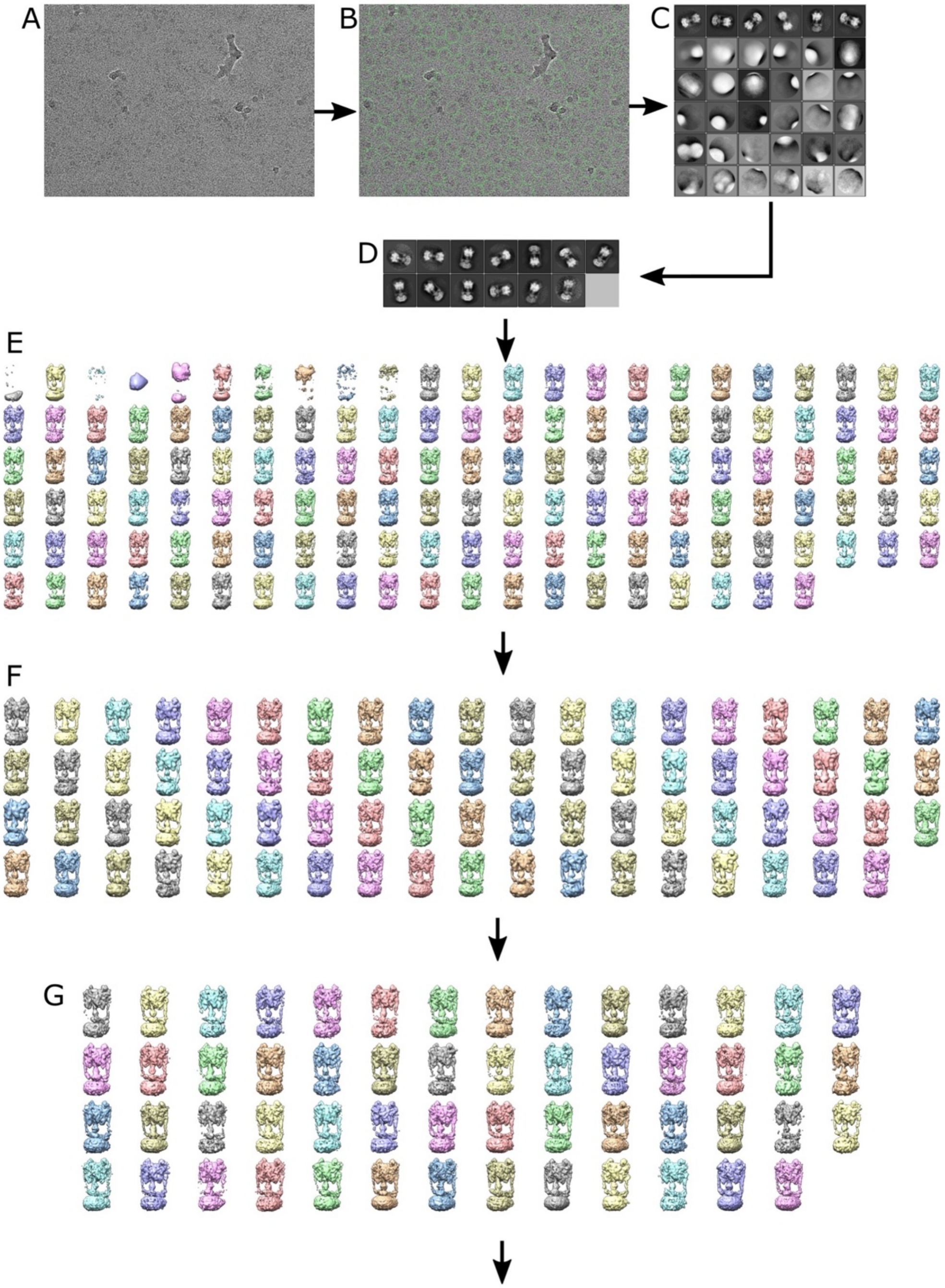


Continues on next page

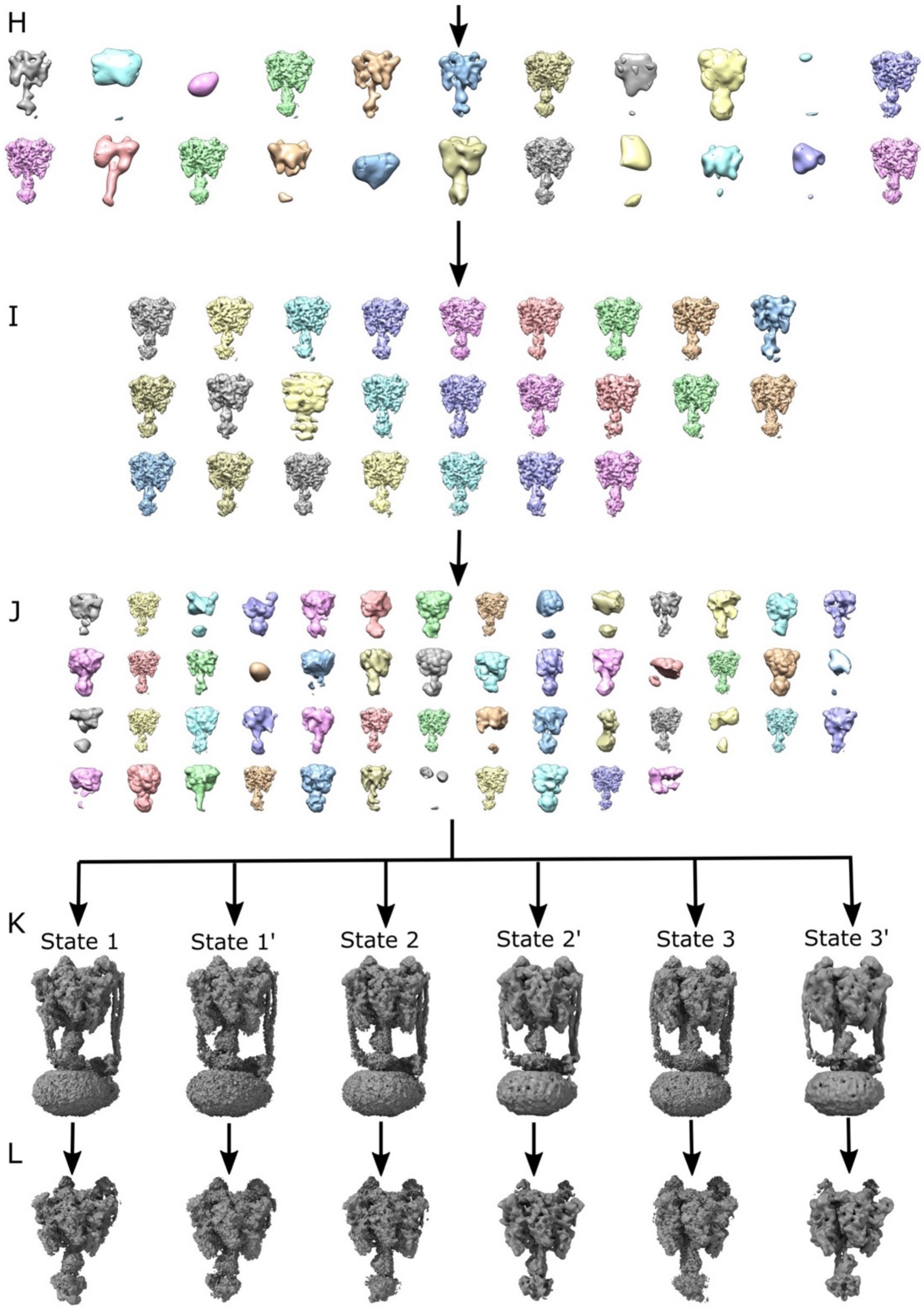
H
I
J
K
State 1
State 1'
State 2
State 2'
State 3
State 3'
L

**Figure S1.** Flowchart showing the processing pathway. A) A representative micrograph. B) The same micrograph with autopicked particles circled in green. C) Classes from a 2D classification of a subset of the picks. D) After recombining all selected 2D classes from the subset runs, a second round of 2D classification was carried out and all classes recognisable as V-ATPase were selected. E) All 3D classes from individual rounds of 3D classification; all classes with a complete $V_o$ domain and rotor were selected. F) All unbroken classes were combined into a single 3D refinement, which was then classified with alignment disabled, classes with features missing were discarded. G) Further 3D classification with alignment disabled. H) A soft mask on the $V_1$ domain was created, and 3D classification carried out, with high-occupancy and high-resolution classes selected. I) After grouping by F-subunit orientation, further 3D classification with alignment disabled was carried out. J) Further selection by F-subunit orientation of classes. 3D classification with alignment disabled was used with each independent orientation of the F-subunit (all classes shown) and was used for a final check. A few classes contained just one or two particles were discarded. K) After generation of a reference with "relion_reconstruct" for each orientation, the complex of each state was passed to 3D refinement and underwent cycles of 3D refinement and CTF refinement until no further resolution improvement was evident. L) Using the $V_1$-focussed mask, $V_1$ domain refinements were carried out on the final particle sets for each orientation. FSC curves of the respective reconstructions are shown in Fig. S4.

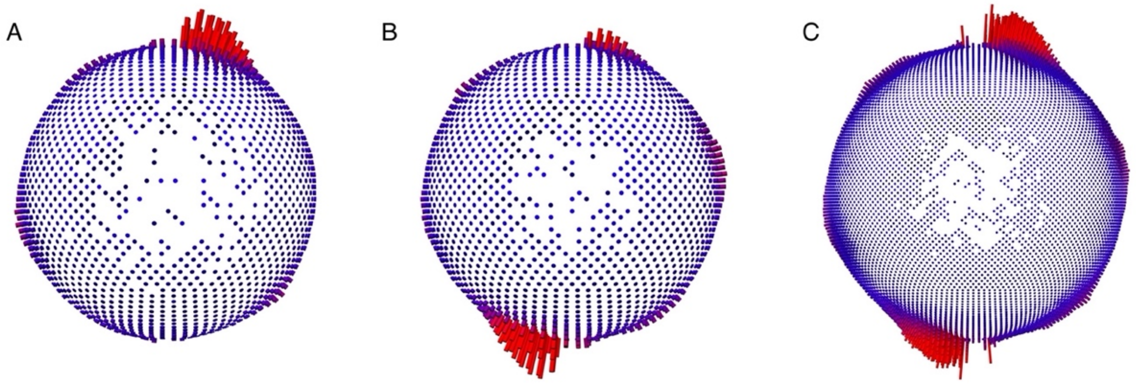


**Figure S2.** Angular assignments showing the orientation preference for the detergent solubilised EhV-ATPase. A) Angles for a 3D refinement from the first dataset. B) Angles from the second and third datasets. C) Combining all datasets permitted the refinements to converge at a finer angular sampling.

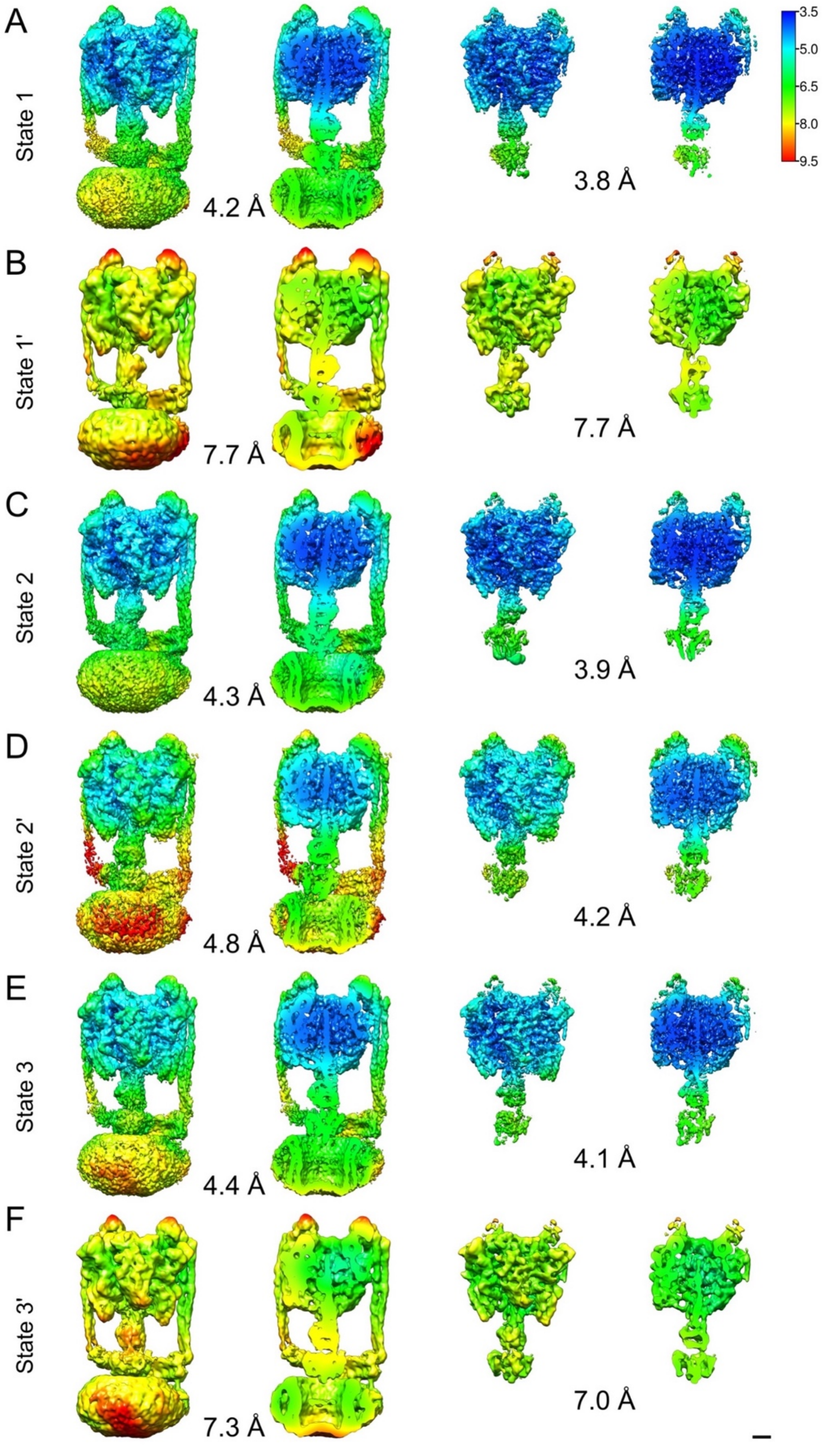
A
State 1
4.2 Å
3.8 Å
3.5
5.0
6.5
8.0
9.5
B
State 1'
7.7 Å
7.7 Å
C
State 2
4.3 Å
3.9 Å
D
State 2'
4.8 Å
4.2 Å
E
State 3
4.4 Å
4.1 Å
F
State 3'
7.3 Å
7.0 Å

55

**Figure S3.** Reconstructions of the six states of EhV-ATPase, filtered and coloured by local resolution as calculated with the local resolution function of RELION 3.1. A) State 1 showing the whole map, the whole map sliced vertically to allow visualisation of internal density and resolution, the $V_1$ focussed refinement, and the $V_1$ focussed refinement sliced vertically to allow visualisation of internal density and resolution. The maps are displayed in the same way as State 1 (A) for State 1' (B), Stare 2 (C), State 2' (D), State 3 (D), and State 3' (F), respectively. Scale bar equals 2 nm.

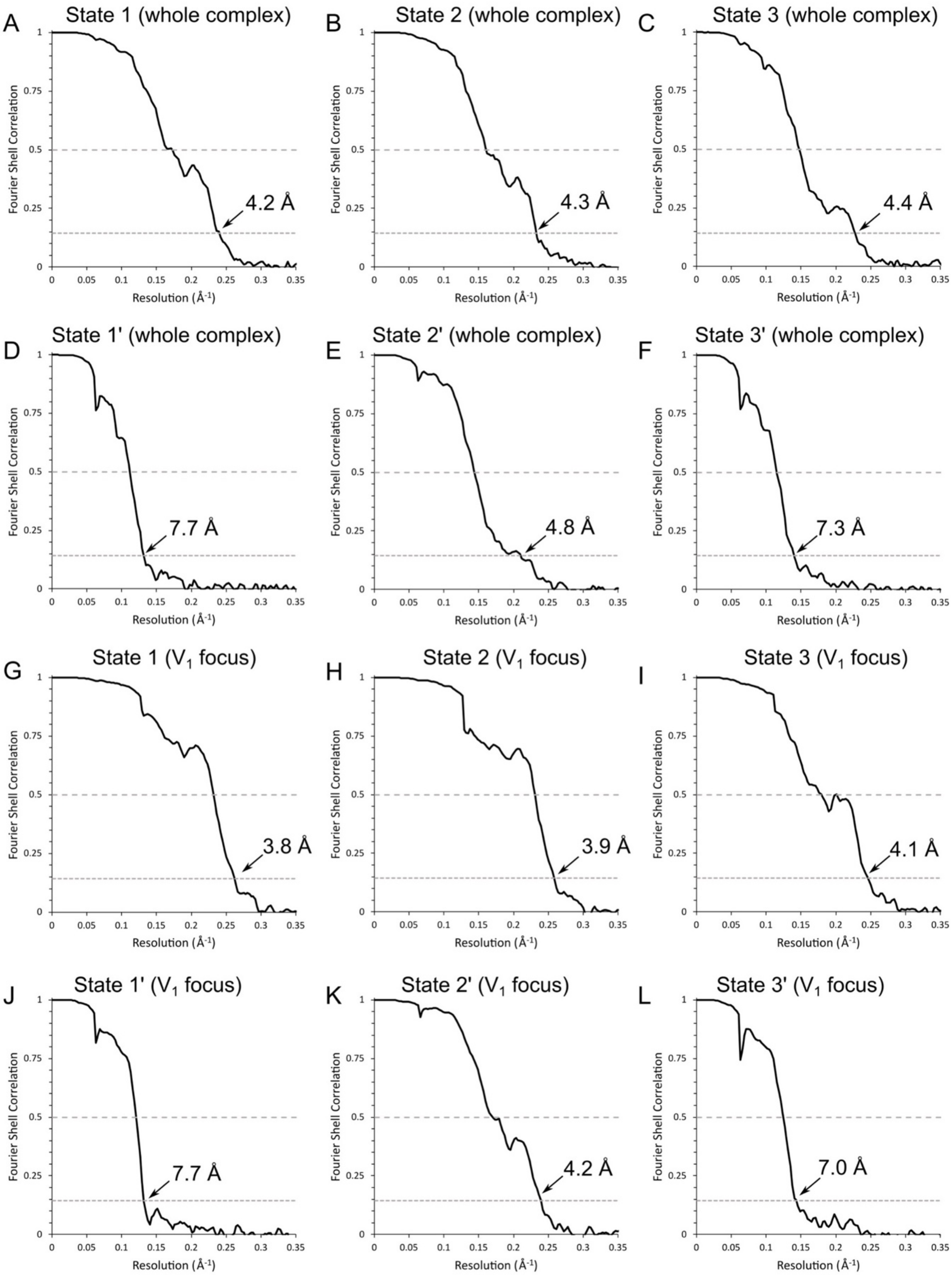
A
State 1 (whole complex)
4.2 Å
B
State 2 (whole complex)
4.3 Å
C
State 3 (whole complex)
4.4 Å
D
State 1' (whole complex)
7.7 Å
E
State 2' (whole complex)
4.8 Å
F
State 3' (whole complex)
7.3 Å
G
State 1 ($V_1$ focus)
3.8 Å
H
State 2 ($V_1$ focus)
3.9 Å
I
State 3 ($V_1$ focus)
4.1 Å
J
State 1' ($V_1$ focus)
7.7 Å
K
State 2' ($V_1$ focus)
4.2 Å
L
State 3' ($V_1$ focus)
7.0 Å
Fourier Shell Correlation
Resolution ($Å^{-1}$)
0
0.25
0.5
0.75
1
0.05
0.1
0.15
0.2
0.25
0.3
0.35

**Figure S4.** Fourier Shell Correlation curves of the twelve reconstructions. A) Whole complex EhV-ATPase at State 1 at 4.2 Å. B) Whole complex EhV-ATPase at State 2 at 4.3 Å. C) Whole complex EhV-ATPase at State 3 at 4.4 Å. D) Whole complex EhV-ATPase at State 1' at 7.7 Å. E) Whole complex EhV-ATPase at State 2' at 4.8 Å. F) Whole complex EhV-ATPase at State 3' at 7.3 Å. G) $V_1$ domain EhV-ATPase at State 1 at 3.8 Å. H) $V_1$ domain EhV-ATPase at State 2 at 3.9 Å. I) $V_1$ domain EhV-ATPase at State 3 at 4.1 Å. J) $V_1$ domain EhV-ATPase at State 1' at 7.7 Å. K) $V_1$ domain EhV-ATPase at State 2' at 4.2 Å. L) $V_1$ domain EhV-ATPase at State 3' at 7.0 Å. All resolutions reported at FSC=0.143.

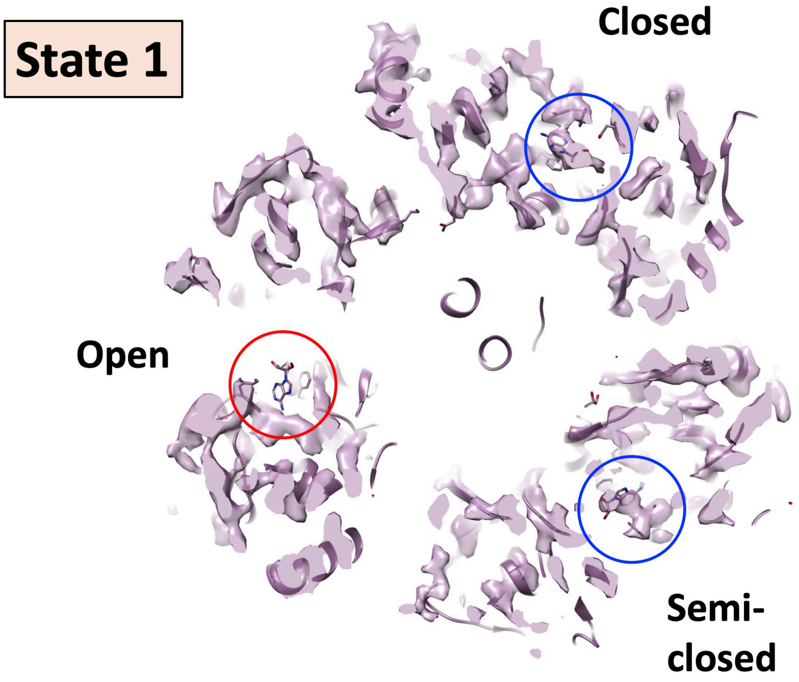


**Figure S5.** The ATP binding pocket densities in EhV-ATPase State 1. The figure is magnified from Fig. 6. The map is sliced through the density map at the level of the ATP binding pocket, and with PDBID:5KNC fitted to check whether density for a bound nucleotide is present or not. Blue circles indicate the presence of density corresponding to the bound nucleotide in the fitted PDB, red circles indicate missing density (only nucleotide from PDB is visible).

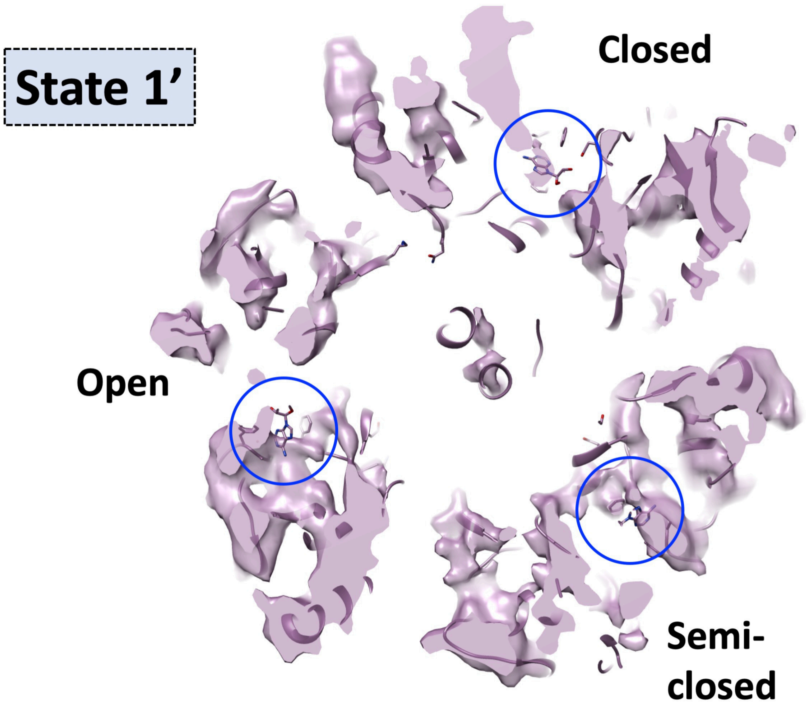


**Figure S6.** The ATP binding pocket densities in EhV-ATPase State 1’. The figure is magnified from Fig. 6. The map is sliced through the density map at the level of the ATP binding pocket, and with PDBID:5KNC fitted to check whether density for a bound nucleotide is present or not. Blue circles indicate the presence of density corresponding to the bound nucleotide in the fitted PDB.

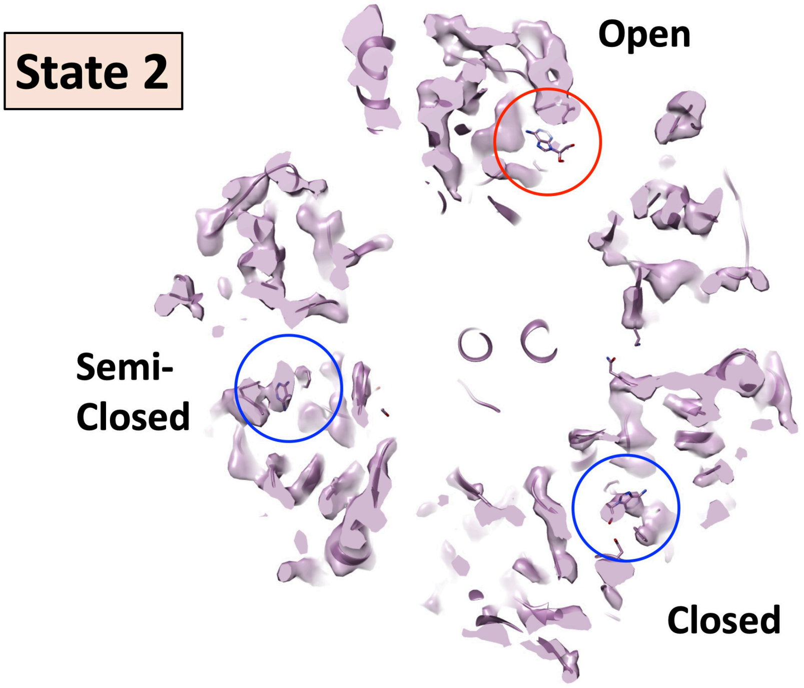


**Figure S7.** The ATP binding pocket densities in EhV-ATPase State 2. The figure is magnified from Fig. 6. The map is sliced through the density map at the level of the ATP binding pocket, and with PDBID:5KNC fitted to check whether density for a bound nucleotide is present or not. Blue circles indicate the presence of density corresponding to the bound nucleotide in the fitted PDB, red circles indicate missing density (only nucleotide from PDB is visible).

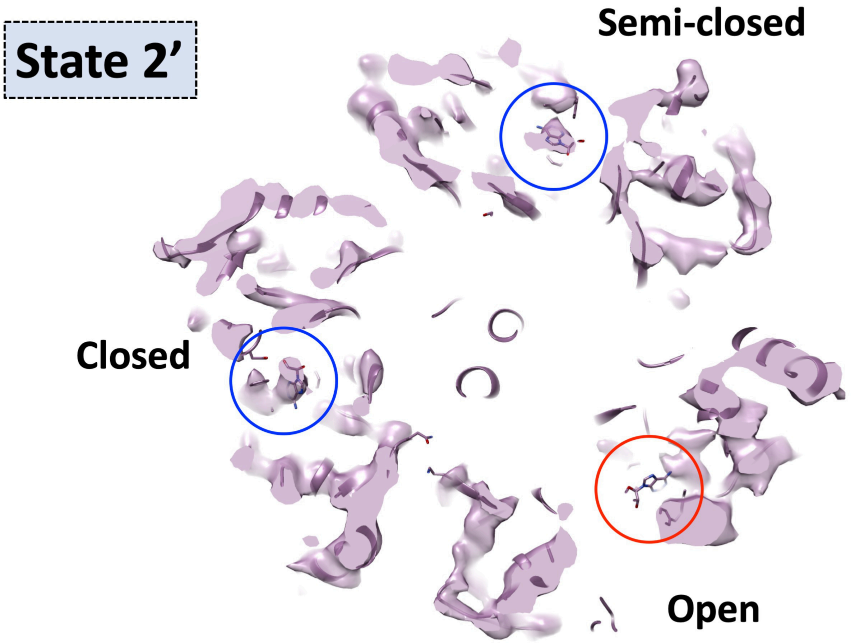


**Figure S8.** The ATP binding pocket densities in EhV-ATPase State 2’. The figure is magnified from Fig. 6. The map is sliced through the density map at the level of the ATP binding pocket, and with PDBID:5KNC fitted to check whether density for a bound nucleotide is present or not. Blue circles indicate the presence of density corresponding to the bound nucleotide in the fitted PDB, red circles indicate missing density (only nucleotide from PDB is visible).

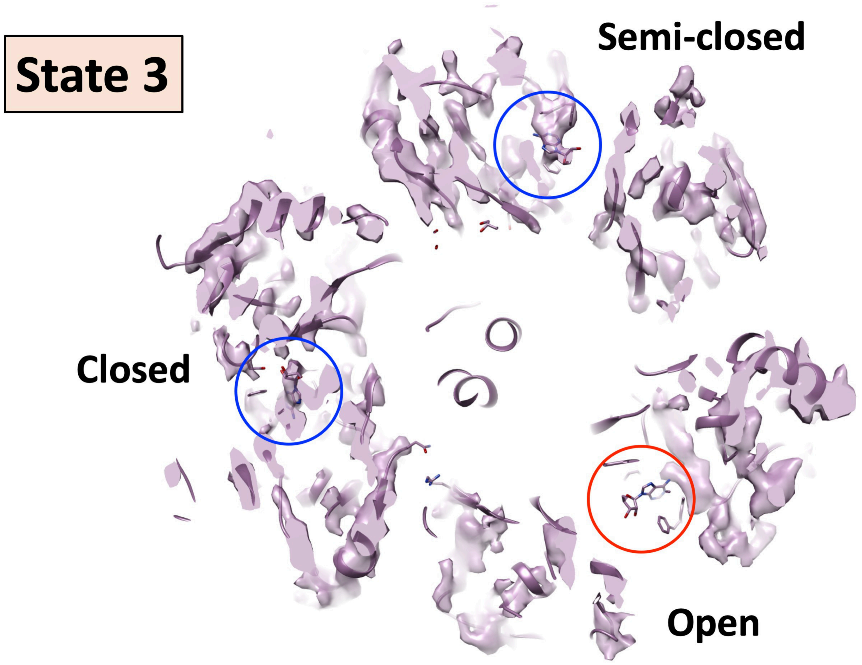


**Figure S9.** The ATP binding pocket densities in EhV-ATPase State 3. The figure is magnified from Fig. 6. The map is sliced through the density map at the level of the ATP binding pocket, and with PDBID:5KNC fitted to check whether density for a bound nucleotide is present or not. Blue circles indicate the presence of density corresponding to the bound nucleotide in the fitted PDB, red circles indicate missing density (only nucleotide from PDB is visible).

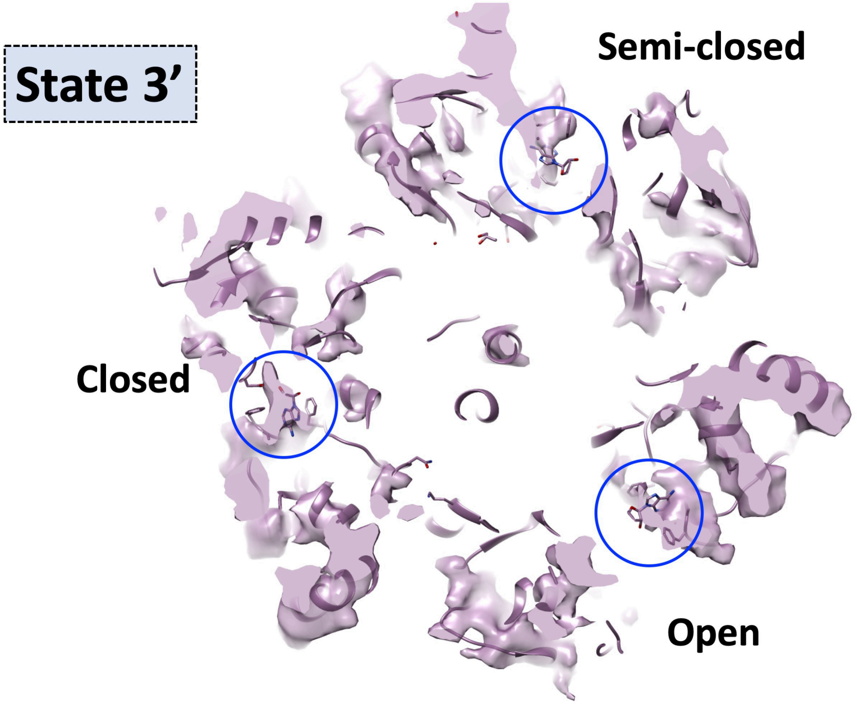


**Figure S10.** The ATP binding pocket densities in EhV-ATPase State 3’. The figure is magnified from Fig. 6. The map is sliced through the density map at the level of the ATP binding pocket, and with PDBID:5KNC fitted to check whether density for a bound nucleotide is present or not. Blue circles indicate the presence of density corresponding to the bound nucleotide in the fitted PDB.
